# Vorapaxar and aripiprazole suppress hepatitis B virus replication through distinct host signaling pathways

**DOI:** 10.64898/2026.08.05.743121

**Authors:** Atsuya Yamashita, Hirotake Kasai, Haruyo Aoyagi, Kousho Wakae, Kaoru Kobayashi, Atsushi Miyajima, Yuichiro Higuchi, Hiroshi Suemizu, Ryo Fukushima, Masanori Isogawa, Takaji Wakita, Hideki Aizaki, Kohji Moriishi

**Affiliations:** Department of Microbiology, Faculty of Medicine, Graduate Faculty of Interdisciplinary Research, University of Yamanashi, Yamanashi 409-3898, Japan; Department of Virology II, National Institute of Infectious Diseases, Japan Institute for Health Security, Tokyo 162-8640, Japan; Department of Biopharmaceutics, Graduate School of Clinical Pharmacy, Meiji Pharmaceutical University, Tokyo, Japan; Central Institute for Experimental Medicine and Life Science, Kawasaki, Japan; Japan Institute for Health Security, Tokyo, Japan; Division of Hepatitis Virology, Institute for Genetic Medicine, Hokkaido University, Hokkaido 060-0815, Japan; Center for Life Science Research, University of Yamanashi, Chuo, Yamanashi, Japan

**Author notes:** Corresponding author: Kohji Moriishi, D.V.M., Ph.D., **E-mail:**. The authors contributed equally (Atsuya Yamashita, Hirotake Kasai).

## Abstract

**Background & Aims:** Current nucleos(t)ide analogs efficiently suppress hepatitis B virus (HBV) replication but have limited effects on viral transcription from covalently closed circular DNA (cccDNA) and integrated HBV DNA. We aimed to identify clinically applicable compounds that directly inhibit HBV transcription by screening FDA-approved drugs.

**Approach & Results:** Screening of 1,470 FDA-approved compounds using an HBV enhancer I/X promoter reporter system identified vorapaxar and aripiprazole as potent inhibitors of viral promoter activity. Both compounds suppressed HBV replication in HBV-producing cells, HBV-infected HepG2-hNTCP cells, and primary human hepatocytes. Aripiprazole reduced hepatocyte nuclear factor 4α (HNF4α) protein levels through an ERK/JNK-dependent pathway and inhibited HBV core promoter activity, whereas vorapaxar acted independently of HNF4α. Both compounds suppressed enhancer I/X promoter activity through inhibition of STAT3 signaling. Vorapaxar inhibited PAR-1-mediated SRC, EGFR, and STAT3 activation, while aripiprazole suppressed SRC–STAT3 signaling independently of EGFR. PAR-1 activation enhanced HBV transcription, whereas PAR-1 knockdown reduced promoter activity and viral RNA expression. Both compounds also reduced HBV replication in human liver chimeric mice at clinically relevant exposure levels without apparent severe toxicity.

**Conclusions:** Vorapaxar and aripiprazole suppress HBV transcription and replication through distinct host signaling pathways. These findings identify PAR-1–STAT3 signaling as a previously unrecognized regulator of HBV transcription and suggest that host-targeting approaches may complement current therapies by suppressing viral gene expression from both cccDNA and integrated HBV DNA.

**Impact and implications:** Current nucleos(t)ide analogues effectively suppress HBV reverse transcription but have limited effects on viral transcription from cccDNA and integrated HBV DNA, highlighting the need for therapies targeting viral gene expression. We identify PAR-1– STAT3 signaling as a previously unrecognized regulator of HBV transcription and demonstrate that two clinically approved drugs, vorapaxar and aripiprazole, suppress HBV replication through distinct host signaling pathways. These findings are relevant to researchers developing host-targeting antivirals and to clinicians seeking complementary therapeutic strategies beyond current nucleos(t)ide analogue therapy. Although further clinical validation and combination studies are required, our results provide a rationale for repurposing approved drugs and for developing transcription-targeting therapies that may complement existing treatments for chronic hepatitis B.

**Highlights:**

- Vorapaxar and aripiprazole suppress HBV through distinct host pathways.
- Both drugs inhibit HBV replication in vitro and in humanized liver mice.
- PAR-1 inhibition reduces HBV transcription by blocking SRC/EGFR/STAT3 signaling.
- PAR-1–STAT3 signaling is a novel regulator of HBV transcription.
- Host-targeting antiviral therapy complements current HBV treatment.

## Introduction

Hepatitis B virus (HBV) chronically infects approximately 296 million people worldwide and remains a major cause of liver cirrhosis and hepatocellular carcinoma despite the availability of effective vaccines and antiviral therapies ^1^. Current nucleos(t)ide analogue therapies efficiently suppress viral reverse transcription and reduce serum HBV DNA levels; however, they have limited effects on the transcriptional activity of covalently closed circular DNA (cccDNA), the persistent viral minichromosome that serves as the template for viral RNA synthesis. In addition, HBV transcripts can be produced from HBV DNA integrated into the host genome, which is not eliminated by current antiviral therapies. Consequently, viral proteins, including hepatitis B surface antigen (HBsAg), often continue to be expressed despite long-term treatment, highlighting the need for therapeutic approaches that directly target HBV transcription.

HBV transcription is regulated by multiple viral promoters and enhancer elements that recruit host transcription factors ^2^. Among these regulatory regions, the enhancer I/X promoter (EnhI/Xp) and the core promoter (Cp), which comprises the core upstream regulatory sequence (CURS) and the basic core promoter (BCP), play central roles in viral gene expression and replication. EnhI/Xp regulates transcription of the HBx gene, which is essential for HBV replication and contributes to the activity of other viral promoters ^3^, whereas CURS-BCP directs the synthesis of precore RNA and pregenomic RNA, both of which are indispensable for viral replication and the production of infectious viral particles. The activities of these regulatory regions are controlled by several liver-enriched and signaling-responsive transcription factors, including hepatocyte nuclear factor 4α (HNF4α) ^4^, FOXA proteins (formerly HNF3 proteins) ^5^, SOX6 ^6^, and signal transducer and activator of transcription 3 (STAT3) ^7^. Previous studies have demonstrated that HNF4α positively regulates CURS-BCP activity and HBV replication ^8,9^, whereas STAT3 cooperates with FOXA proteins to activate enhancer I-mediated transcription ^7,10^. In addition, EGFR signaling has been reported to promote HBV replication through STAT3 phosphorylation ^10^. HBV infection itself also induces STAT3 phosphorylation, thereby enhancing viral replication while protecting infected hepatocytes from HBV-induced apoptosis ^11^. Collectively, these findings suggest that modulation of host signaling pathways regulating key transcription factors, particularly STAT3, represents an attractive strategy for suppressing HBV replication.

Drug repurposing has emerged as an efficient strategy for identifying novel antiviral agents because the pharmacological and safety profiles of approved drugs are already well established. In the present study, we performed a high-throughput screening of FDA-approved compounds using an EnhI/Xp-driven reporter system and identified vorapaxar and aripiprazole as potent inhibitors of HBV transcription. Vorapaxar is a first-in-class, potent, reversible antagonist of protease-activated receptor-1 (PAR-1) ^12,13^, whereas aripiprazole is a third-generation antipsychotic with a favorable safety profile and a relatively low risk of tardive dyskinesia compared with other antipsychotic agents ^14,15^. We subsequently evaluated their antiviral activities, identified the viral regulatory regions responsible for their inhibitory effects, and investigated the underlying molecular mechanisms both in vitro and in vivo. Although both compounds suppressed HBV transcription, whether they act through a common or distinct molecular mechanism remained unknown.

## Results

### Screening for compounds targeting enhancer I-X promoter (EnhI/Xp) activity

We previously established a Huh7-derived reporter cell line expressing firefly luciferase under the control of the HBV enhancer I/X promoter (Huh7GL4.18EnhIX-Pro_AeUS cells) ^8,9^ (Supplementary Figure S1A). This reporter cell line, hereafter referred to as the EnhI/Xp reporter cell line, was used to screen an FDA-approved drug library for inhibitors of HBV transcription. Cells were treated with each compound at a final concentration of 20 µM for 48 h, after which luciferase activity and cell viability were evaluated (Supplementary Figure S1B). Compounds reducing luciferase activity to <35% while maintaining >80% cell viability were defined as primary hits. Twenty-two compounds met these criteria (Supplementary Figure S1B) and were subjected to secondary screening to determine their EC_50_ and CC_50_ values using the EnhI/Xp reporter cell line (Figure 1C). Dose-response analyses identified four compounds— aripiprazole, cariprazine, fosaprepitant dimeglumine, and vorapaxar—with EC_50_ values below 13.7 µM and CC_50_ values above 31.3 µM (Supplementary Figure S1B,D).

**Figure 1.**
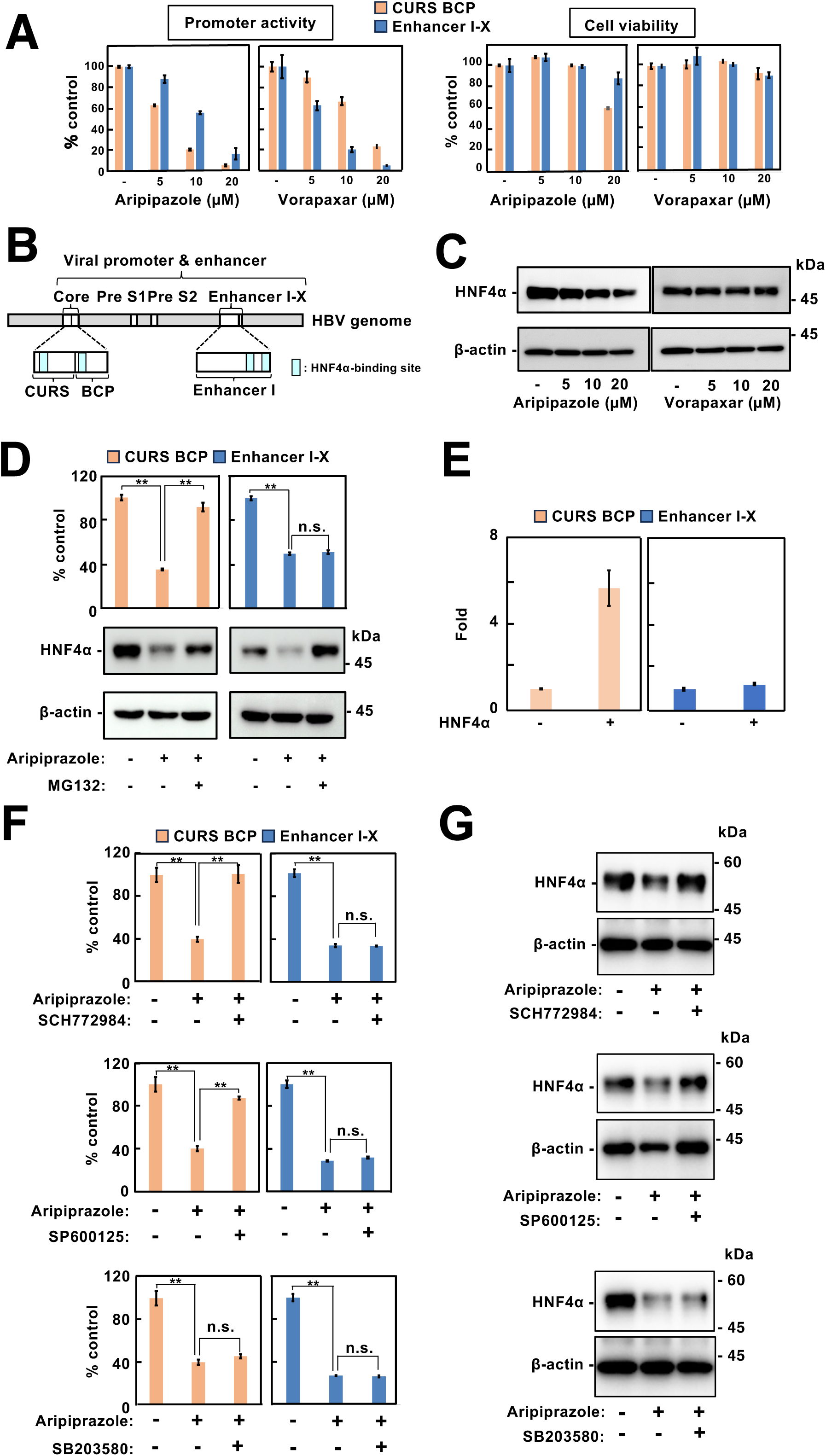
Distinct mechanisms of HBV promoter suppression by aripiprazole and vorapaxar. (A) Effects of aripiprazole and vorapaxar on viral promoter activity. EnhI/Xp- or CURS/BCP-reporter cells were treated with various concentrations of aripiprazole or vorapaxar. Luciferase activity and cell viability were measured as described in the Materials and Methods. (B) Schematic representation of EnhI/Xp and CURS/BCP. Putative HNF4α-binding sites are indicated in EnhI/Xp and CURS/BCP regions. (C) Effects of aripiprazole and vorapaxar on HNF4α protein levels. Reporter cells were treated with aripiprazole or vorapaxar and harvested 8 h after treatment. Cell lysates were subjected to western blot analysis to evaluate HNF4α protein levels. (D) Involvement of the proteasome in the regulation of HNF4α protein levels and reporter activity. Reporter cells were treated with aripiprazole in the presence or absence of the proteasome inhibitor MG132 and harvested 8 h after treatment. Cell lysates were subjected to western blot analysis to evaluate HNF4α protein levels. (E) Effects of HNF4α on HBV promoter activities. HEK293T cells were co-transfected with luciferase reporter plasmids driven by EnhI/Xp or CURS/BCP together with an HNF4α expression plasmid. Firefly luciferase (Fluc) and Renilla luciferase (Rluc) activities were measured, and Fluc activity was normalized to Rluc activity. (F, G) Effects of MAPK inhibitors on viral promoter activity and HNF4α protein levels. Reporter cells were treated with aripiprazole in the presence or absence of MAPK inhibitors (ERK inhibitor, SCH772984; JNK inhibitor, SP600125; p38 inhibitor, SB203580) and harvested 8 h after treatment. Cell lysates were subjected to luciferase assays (F) and western blot analysis (G). Data are presented as the mean ± standard deviation (SD) from three independent experiments.

The antiviral activities of these four compounds were subsequently evaluated in multiple HBV infection models. In HepG2.2.15.7 cells, all compounds exhibited EC_50_ values below 0.5 µM with selectivity indices (SI) greater than 50 (Table 1). In HBV-infected HepG2-hNTCP-C4 cells, EC_50_ values ranged from 0.5 to 4.2 µM, with SI values ranging from 62.5 to >1000 (Table 2). All four compounds also inhibited HBV replication in primary human hepatocytes (Supplementary Figure S2A) and reduced cccDNA production in Hep38.7-Tet cells (Supplementary Figure S2B). These findings demonstrate that aripiprazole, cariprazine, fosaprepitant dimeglumine, and vorapaxar possess potent anti-HBV activity in vitro. Pharmacokinetic simulations predicted that cariprazine and fosaprepitant dimeglumine would not accumulate sufficiently in the murine liver to achieve therapeutic concentrations (data not shown). Therefore, subsequent analyses focused on vorapaxar and aripiprazole.

**Table 1.**
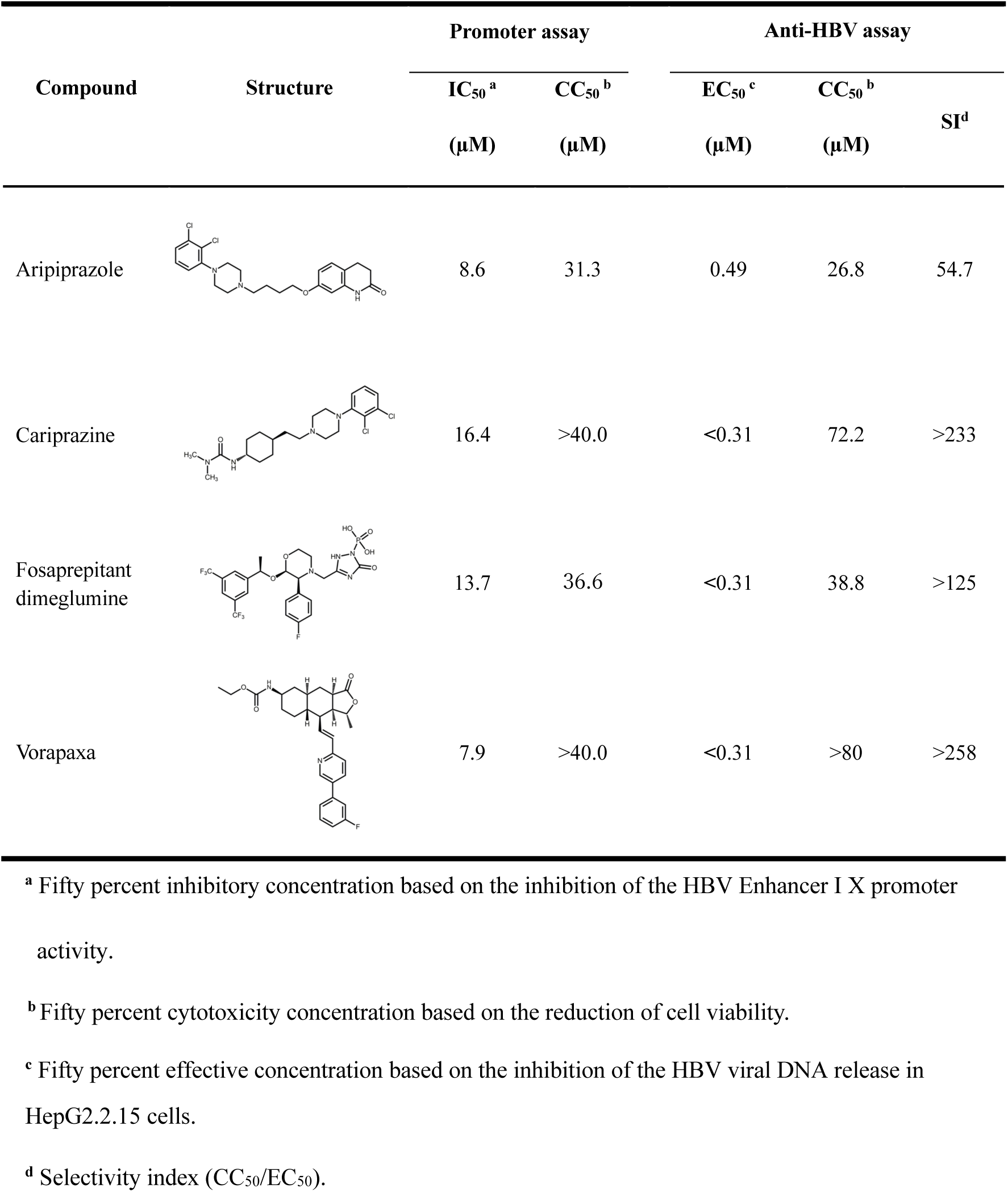
Effects of 2nd hit compounds on HBV promoter activity and anti HBV activity.

| Compound | Structure | Promoter assay |  | Anti-HBV assay |  | SI<sup>d</sup> |
| --- | --- | --- | --- | --- | --- | --- |
|  |  | IC<sub>50</sub><sup>a</sup> | CC<sub>50</sub><sup>b</sup> | EC<sub>50</sub><sup>c</sup> | CC<sub>50</sub><sup>b</sup> |  |
| | | (μM) | (μM) | (μM) | (μM) | |
| Aripiprazole |  | 8.6 | 31.3 | 0.49 | 26.8 | 54.7 |
| Cariprazine |  | 16.4 | >40.0 | <0.31 | 72.2 | >233 |
| Fosaprepitant dimeglumine |  | 13.7 | 36.6 | <0.31 | 38.8 | >125 |
| Vorapaxa |  | 7.9 | >40.0 | <0.31 | >80 | >258 |

<sup>a</sup> Fifty percent inhibitory concentration based on the inhibition of the HBV Enhancer I X promoter activity.
<sup>b</sup> Fifty percent cytotoxicity concentration based on the reduction of cell viability.
<sup>c</sup> Fifty percent effective concentration based on the inhibition of the HBV viral DNA release in HepG2.2.15 cells.
<sup>d</sup> Selectivity index (CC<sub>50</sub>/EC<sub>50</sub>).

**Table 2.** Anti-HBV activities of 2nd hit compounds in cell line-based HBV infection system.

| Name | EC<sub>50</sub><sup>a</sup><br>(μM) | CC<sub>50</sub><sup>b</sup><br>(μM) | SI<sup>c</sup> |
| --- | --- | --- | --- |
| Aripiprazole | 4.2 ± 4.3 | 77.5 ± 13.8 | 62.5 |
| Cariprazine | 0.9 ± 0.4 | 260.0 ± 27.4 | 332.6 |
| Fosaprepitant<br>dimeglumine | 2.2 ± 2.2 | 164.0 ± 48.9 | 195.7 |
| Vorapaxar | 0.5 ± 0.6 | > 500 | > 1000 |

<sup>a</sup> Fifty percent effective concentration based on the inhibition of the HBV viral DNA release.
<sup>b</sup> Fifty percent cytotoxicity concentration based on the reduction of cell viability.
<sup>c</sup> Selectivity index (CC<sub>50</sub>/EC<sub>50</sub>).

### Effect of vorapaxar and aripiprazole on HBV core promoter activity

To determine whether vorapaxar and aripiprazole also affect HBV core promoter (Cp) activity, we used a Huh7-derived reporter cell line expressing luciferase under the control of the core upstream regulatory sequence (CURS) and basal core promoter (BCP) (Huh7GL4.18CURS_BC_AeUS cells) ^9^, hereafter referred to as the Cp reporter cell line. Treatment with either vorapaxar or aripiprazole suppressed Cp activity to a similar extent as observed for EnhI/Xp activity (Figure 1A). The CURS-BCP region contains well-characterized HNF4α-binding sites, whereas two putative HNF4α-binding sites are predicted within enhancer I (Figure 1B). Aripiprazole markedly reduced HNF4α protein levels, whereas vorapaxar had no detectable effect (Figure 1C). MG132 restored HNF4α protein expression in aripiprazole-treated cells and abolished the inhibitory effect of aripiprazole on CURS-BCP activity but not on EnhI/Xp activity (Figure 1D). Consistently, HNF4α overexpression enhanced CURS-BCP activity but had little effect on EnhI/Xp activity (Figure 1E). To investigate the involvement of MAP kinase signaling, cells were treated with inhibitors of ERK (SCH772984), JNK (SP600125), or p38 MAPK (SB203580). Inhibition of ERK or JNK restored both HNF4α protein levels and CURS-BCP activity in aripiprazole-treated cells, whereas inhibition of p38 MAPK had no effect (Figure 1F,G). In contrast, vorapaxar did not alter HNF4α protein abundance under any condition examined. These findings indicate that aripiprazole suppresses HBV core promoter activity through ERK- and JNK-dependent downregulation of HNF4α, whereas vorapaxar acts through an HNF4α-independent mechanism.

### Determination of the antiviral-target regions within EnhI/Xp

To identify the regions within EnhI/Xp responsible for the antiviral effects of vorapaxar and aripiprazole, EnhI/Xp was subdivided into seven regulatory domains (D1–D7) based on predicted transcription factor-binding sites (Supplementary Figure S3). Reporter constructs carrying deletions of individual domains were generated (Figure 2A, left panel). In the absence of compounds, deletion of D3, D4, D5, or D6 significantly reduced promoter activity (Figure 2A, right panel). Reporter constructs containing six tandem repeats of individual domains further demonstrated that D4, D5, and D6 possessed substantially higher enhancer activity than D3 (Figure 2B). Both vorapaxar and aripiprazole markedly inhibited the activities of D4- and D5-driven reporters but had little effect on D6-driven reporters (Figure 2C), indicating that D4 and D5 represent the principal target regions of both compounds. Putative STAT3-binding sites were predicted within both D4 and D5, whereas a putative FOXA-binding site was identified within D5 (Supplementary Figure 3). Based on these findings, we next investigated whether vorapaxar and aripiprazole affect STAT3 signaling.

**Figure 2.**
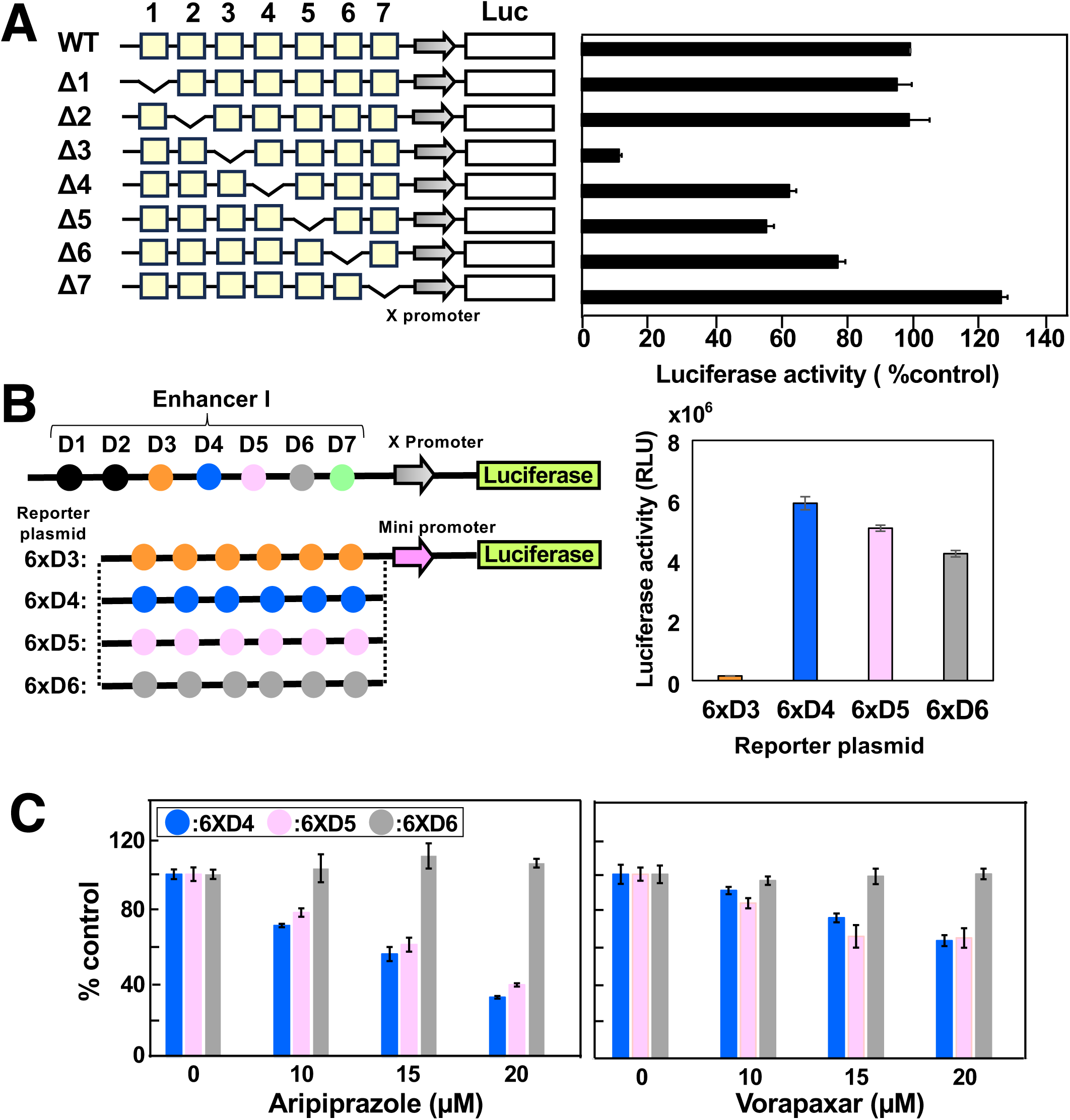
Identification of EnhI/Xp domains mediating basal transcriptional activity and responsiveness to aripiprazole and vorapaxar. For all panels, Fluc activity was normalized to Rluc activity using the pRL-TK control vector. (A) Functional analysis of EnhI/Xp deletion-mutants. Luciferase reporter plasmids containing the wild-type (WT) EnhI/Xp or deletion mutants lacking domains D1–D7 (Δ1 to Δ7) were transfected into Huh7 cells. Luciferase activity was measured and normalized to that of the WT promoter, which was set to 100%. (B) Transcriptional activities of multimerized EnhI/Xp domains. Luciferase reporter plasmids containing six tandem repeats of domains D1–D7 upstream of a minimal promoter were transfected into Huh7 cells. Fluc activity was measured and normalized to Rluc activity. (C) Effects of aripiprazole and vorapaxar on multimerized domain-driven reporter activity. Huh7 cells transfected with luciferase reporter plasmids containing six tandem repeats of domains D4, D5, D6, or D7 were treated with aripiprazole or vorapaxar. Luciferase activity was measured and expressed as a percentage of the corresponding untreated control. (B, C) “6xD3”, “6xD4”, “6xD5” and “6xD6” indicate pGL4.18EnhI-Xpro6xD3, pGL4.18EnhI-Xpro6xD4, pGL4.18EnhI-Xpro6xD5 and pGL4.18EnhI-Xpro6xD6, respectively. Data are presented as the mean ± SD from three independent experiments.

### Effect of vorapaxar and aripiprazole on STAT-3 phosphorylation and upstream signaling molecules

Because D4 and D5 contain putative STAT3-binding sites (Supplementary Figure S3), we examined the phosphorylation status of STAT3 and its upstream signaling molecules following treatment with vorapaxar or aripiprazole. EnhI/Xp reporter cells were seeded at 1 × 10^5^ cells per well and treated with 20 µM vorapaxar or aripiprazole 24 h after seeding. Cell lysates were collected 4 or 8 h after treatment and analyzed by western blotting. Both vorapaxar and aripiprazole reduced phosphorylation of STAT3 and SRC (Figure 3A,B). In contrast, only vorapaxar decreased EGFR phosphorylation, whereas aripiprazole had little effect on EGFR phosphorylation (Figure 3A,B). These findings suggest that the two compounds suppress STAT3 activation through distinct upstream signaling pathways. We therefore next investigated the role of PAR-1 signaling in STAT3 activation and HBV replication.

**Figure 3.**
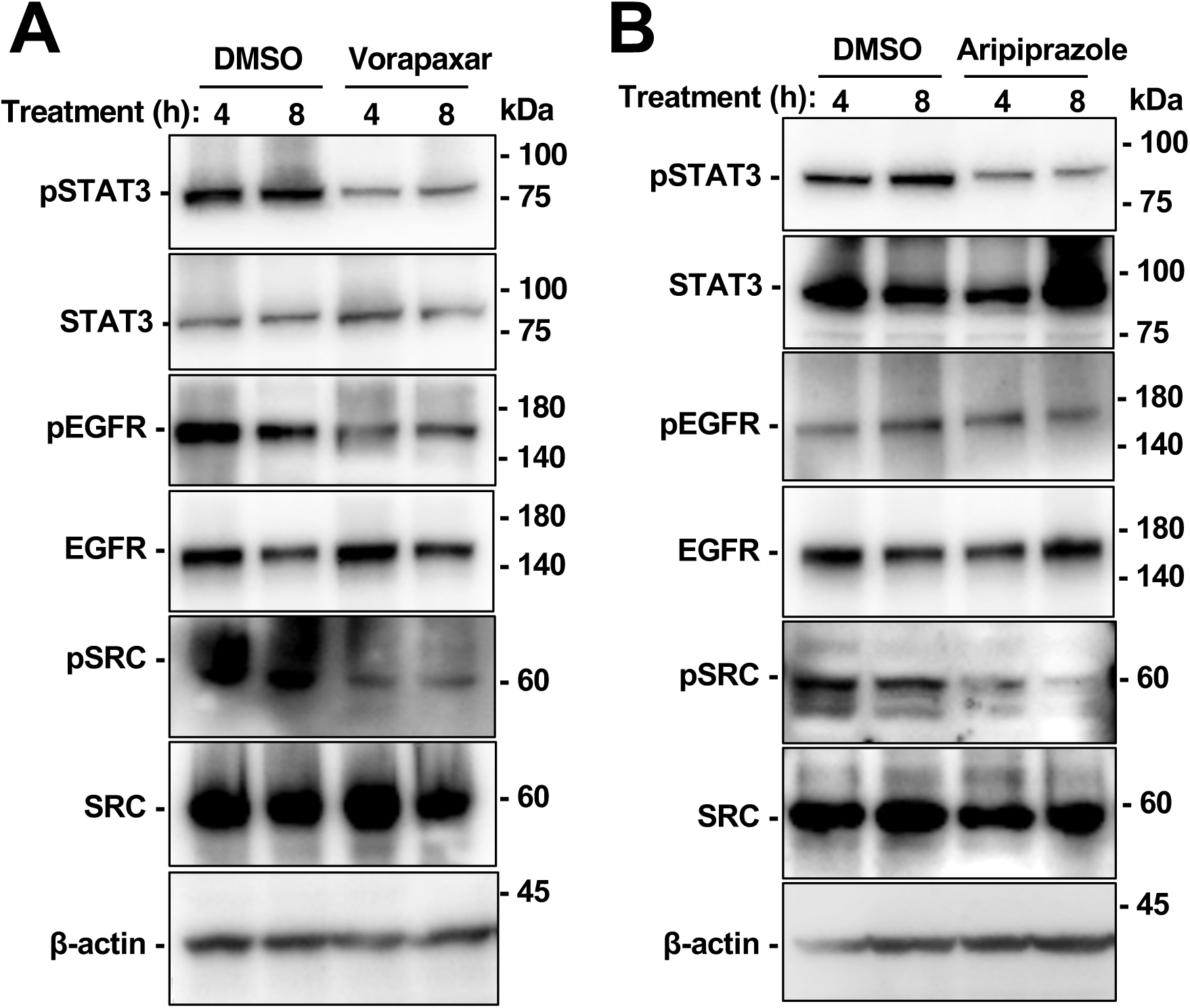
Effects of vorapaxar and aripiprazole on STAT3, EGFR, and c-SRC Phosphorylation. EnhI/Xp reporter cells (1 × 10⁵ cells) were treated with aripiprazole (A), vorapaxar (B) or vehicle control (DMSO). The treated cells were harvested 4h or 8 h after treatment to prepare cell lysates. These cell lysates were subjected into western blotting.

### Role of PAR-1 in STAT3 activation and HBV replication

To confirm PAR-1 expression, total RNA was isolated from Huh7 cells and analyzed by RT-sqPCR using threefold serial dilutions. PAR-1 mRNA was readily detected as a single specific amplicon, and its amplification was proportional to the amount of input RNA, similar to GAPDH (Figure 4A). Knockdown of PAR-1 using siRNA significantly reduced PAR-1 mRNA expression as well as EnhI/Xp and Cp reporter activities (Figure 4B). To determine whether PAR-1 regulates HBV replication in primary human hepatocytes, PXB cells were infected with HBV and transfected with siPAR-1 at 12 days post-infection. PAR-1 knockdown significantly reduced both PAR-1 mRNA expression and total HBV RNA levels (Figure 4C). To further evaluate the effect of PAR-1 activation, EnhI/Xp and Cp reporter cells were treated with the PAR-1-agonist peptide TFLLR-NH2. PAR-1 activation significantly enhanced both EnhI/Xp and Cp promoter activities (Figure 4D) and increased HBV RNA levels in HBV-infected HepG2-hNTCP cells (Figure 4E). Flow cytometric analysis demonstrated that PAR-1 knockdown reduced cell-surface PAR-1 expression (Figure 5A,B). Consistently, PAR-1 knockdown decreased phosphorylation of STAT3 and EGFR (Figure 5C), whereas PAR-1 overexpression increased cell-surface PAR-1 expression together with phosphorylation of SRC and EGFR in a dose-dependent manner (Figure 5D–F). Collectively, these findings indicate that PAR-1 positively regulates HBV promoter activity through activation of the SRC/EGFR/STAT3 signaling pathway.

**Figure 4.**
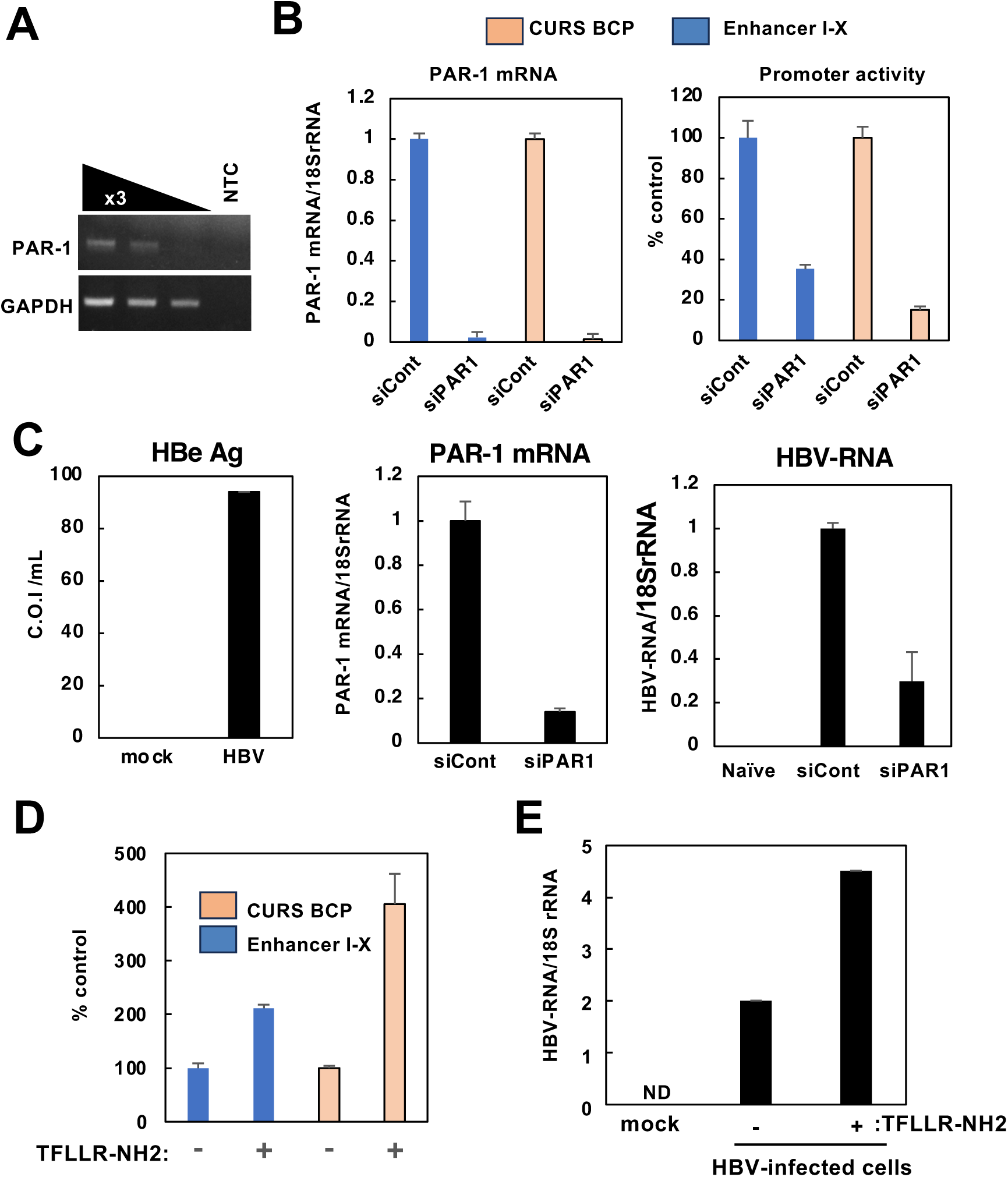
Evaluation of the role of protease-activated receptor 1 (PAR-1), the molecular target of vorapaxar, in HBV transcription. (A) Expression of endogenous PAR-1 in Huh7 cells. Total RNA was isolated from Huh7 cells, from which these reporter cell lines were originated. PAR-1 mRNA levels were assessed by RT-sqPCR using threefold serial dilutions of total RNA. (B) Effects of PAR-1 knockdown on HBV promoter activity. PAR-1 mRNA levels were measured 72 h after transfection with PAR-1-targeting siRNA (siPAR1) in EnhI/Xp reporter cells and CURS/BCP reporter cells (a left panel). Promoter activities were evaluated in parallel (a right panel). (C) Effects of PAR-1 knockdown on HBV transcription in HBV-infected cells. PAR-1-targeting siRNA was introduced into HBV-infected PXB cells at 12 days post-infection (dpi; MOI = 5). Left panel, HBeAg levels in culture supernatants at 12 dpi; center panel, PAR-1 mRNA levels at 48 h after siRNA transfection; right panel, HBV RNA levels at 72 h after siRNA transfection. (D) Effects of PAR-1 activation on HBV promoter activity. EnhI/Xp and CURS/BCP reporter cells were treated with 20 µM of PAR-1-activating peptide TFLLR-NH2, and promoter activities were measured 24 h after treatment. (E) Effects of PAR-1 activation on HBV transcription in infected cells. HepG2-hNTCP-C4 cells were treated with the PAR-1-activating peptide 12 days after HBV infection. Total RNA was harvested 24 h after peptide treatment, and HBV RNA levels were quantified by RT-qPCR.

**Figure 5.**
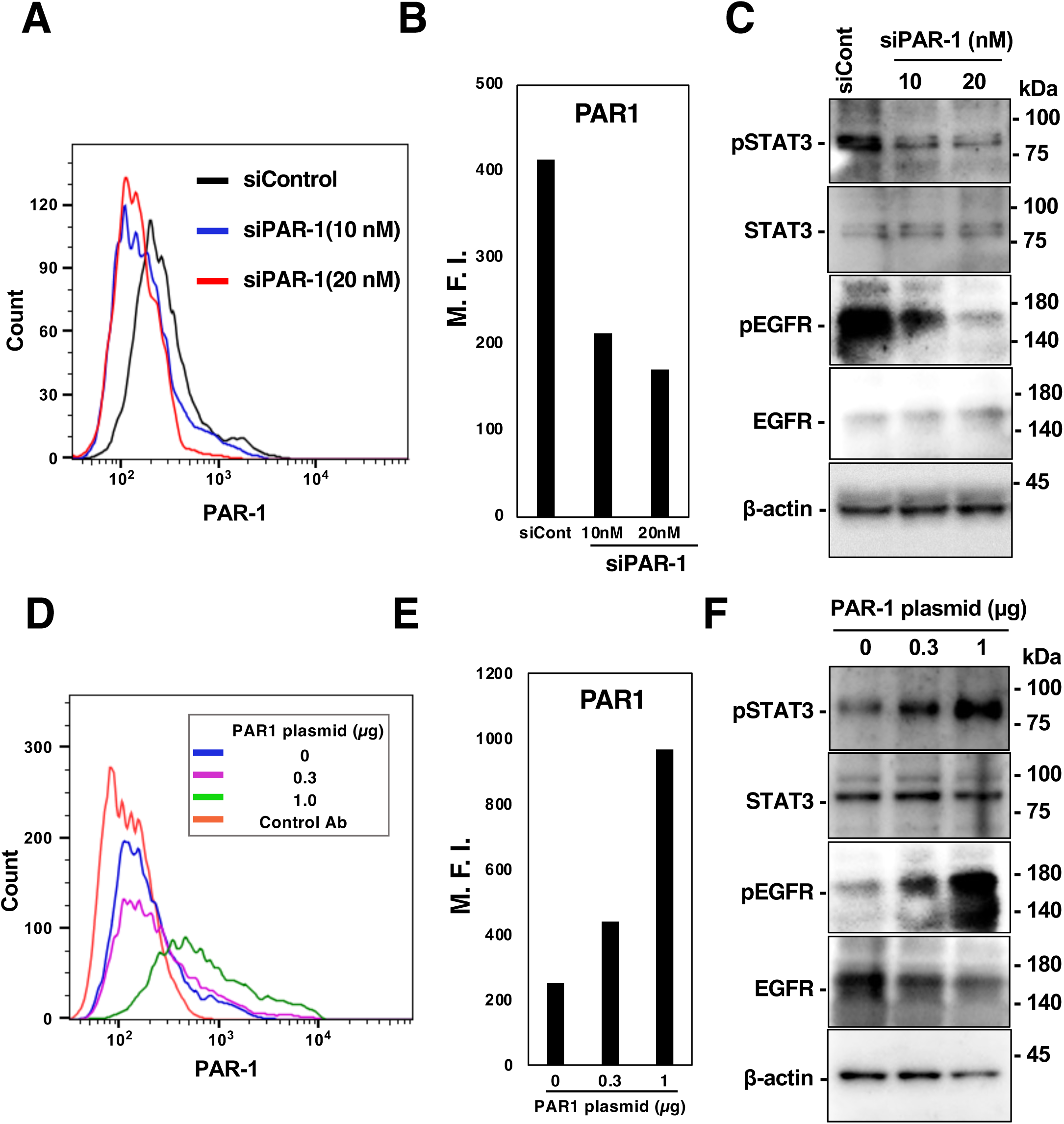
Effects of PAR-1 knockdown or overexpression on STAT3 phosphorylation. (A–C) Effects of PAR-1 knockdown on STAT3 phosphorylation. EnhI/Xp reporter cells were transfected with PAR-1-targeting siRNA (siPAR-1) or control siRNA (siCont) and harvested 48 h after transfection. Cell-surface PAR-1 expression was analyzed by flow cytometry using an anti-PAR-1 antibody. Representative flow cytometry plots are shown (A), and the mean fluorescence intensity (MFI) was quantified (B). Cell lysates were subjected to western blot analysis to evaluate STAT3 phosphorylation and related proteins (C). (D–F) Effects of PAR-1 overexpression on STAT3 phosphorylation. EnhI/Xp reporter cells were transfected with a PAR-1 expression plasmid or an empty vector and harvested 48 h after transfection. To maintain a constant amount of transfected DNA, the amount of PAR-1 expression plasmid (0–1 µg/plate) was adjusted with empty vector. Cell-surface PAR-1 expression was analyzed by flow cytometry using an anti-PAR-1 antibody. Representative flow cytometry plots are shown (D), and the mean fluorescence intensity (MFI) was quantified (E). Cell lysates were subjected to western blot analysis to evaluate STAT3 phosphorylation and related proteins (F).

### Antiviral effects of vorapaxar and aripiprazole in human liver chimeric mice

We next evaluated the anti-HBV activities of vorapaxar and aripiprazole in human liver chimeric mice. Following a single oral administration of vorapaxar (125 mg/kg), plasma and intrahepatic drug concentrations exceeded the in vitro EC_90_ (Supplementary Figure 6A). Because the elimination half-life was not determined in this study, the previously reported value in rats (5.1 h) ^16^ was used for pharmacokinetic simulation. Based on the reported maximum clinical dose of vorapaxar (0.7 mg/kg/day), a dosing regimen of 0.5 mg/kg every 12 h was selected for simulation of repeated administration ^17^. Plasma concentrations following repeated administration (0.5 mg/kg every 12 h) were predicted to remain above the in vitro EC_50_ (Supplementary Figure 6B). All transplanted mice with sufficient human hepatocyte repopulation developed stable HBV viremia after viral inoculation (Supplementary Figure S4A). Oral administration of vorapaxar (0.5 mg/kg every 12 h) was initiated at 32 days post-inoculation, when serum HBV DNA levels had reached a plateau. Vorapaxar treatment significantly reduced serum HBV DNA levels after two weeks compared with the control group (Supplementary Figure S4B and Figure 6C).

**Figure 6.**
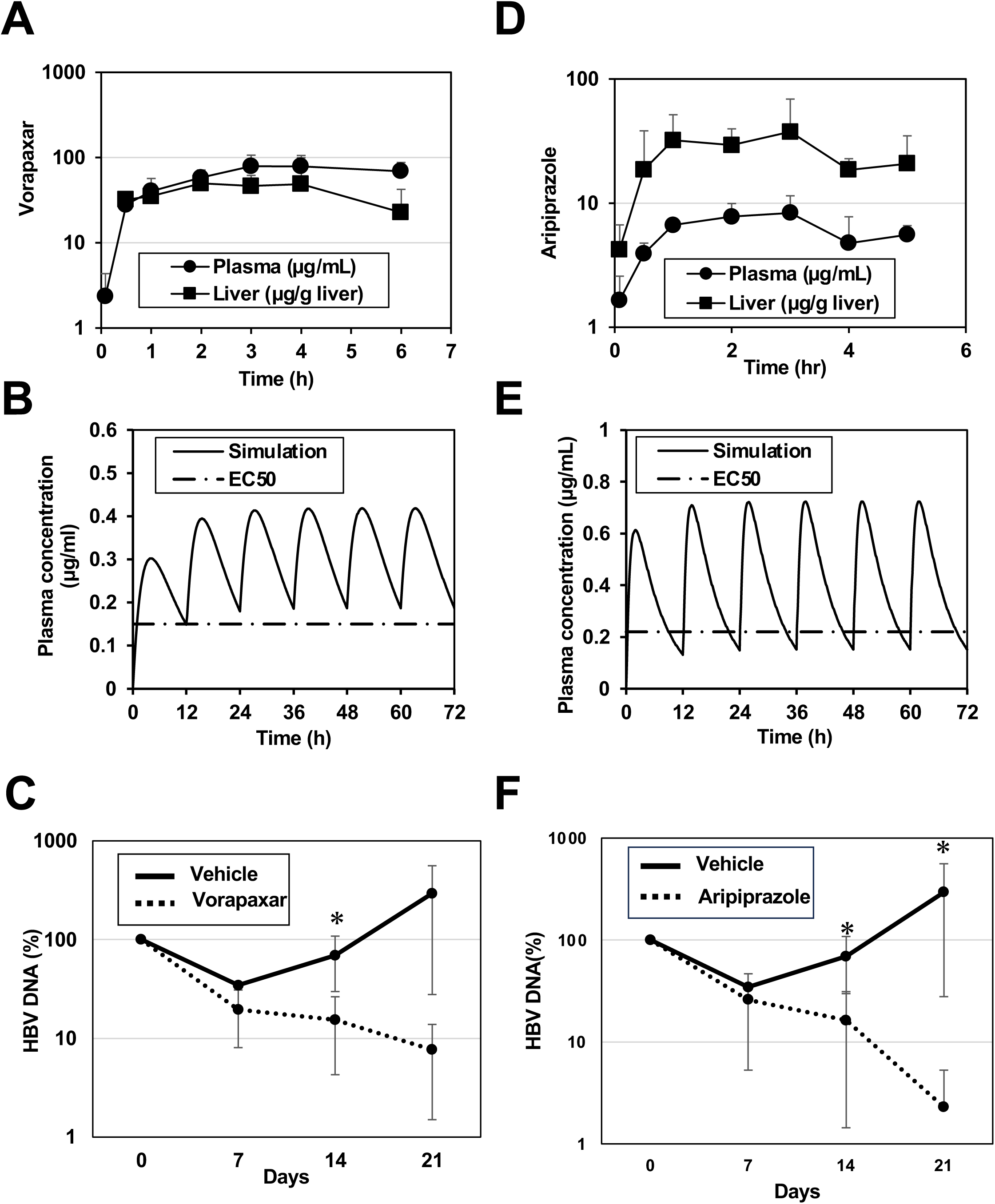
Distinct host-targeting antivirals, vorapaxar and aripiprazole, suppress serum HBV DNA in NOG-TKm30 human liver chimeric mice. Mice with serum cholinesterase levels greater than 50 U/L, indicative of efficient and stable human hepatocyte engraftment, were inoculated with HBV (1 × 10^9^ copies/mouse) via the tail vein on day −32. HBV-infected NOG-TKm30 human liver chimeric mice with stable viremia were orally administered vehicle, vorapaxar, or aripiprazole at a dose of 0.5 mg/kg twice daily for 21 consecutive days. (A) Plasma and intrahepatic concentration profiles of vorapaxar in wild-type mice. Mice (n = 4) were administered 125 mg/kg vorapaxar. Data are expressed as mean ± S.D. (B) Pharmacokinetic simulation of plasma vorapaxar concentrations following multiple oral administrations of 0.5 mg/kg every 12 h. The dotted line indicates the in vitro EC50 of vorapaxar. (C) Time course of serum HBV DNA levels, expressed as a percentage of baseline (Day 0 = 100%). Data are shown as mean ± SEM. Statistical significance was determined using appropriate parametric or non-parametric tests. A p value < 0.05 was considered statistically significant. (D) Plasma and intrahepatic concentration profiles of aripiprazole in wild-type mice. Mice (n = 4) were administered 125 mg/kg aripiprazole. Data are expressed as mean ± S.D. (E) Pharmacokinetic simulation of plasma aripiprazole concentrations following multiple oral administrations of 10 mg/kg every 12 h. The dotted line indicates the in vitro EC50 of aripiprazole. (F) Time course of serum HBV DNA levels, expressed as a percentage of baseline (Day 0 = 100%). Data are shown as mean ± SEM. Statistical significance was determined using appropriate parametric or non-parametric tests. A p value < 0.05 was considered statistically significant.

Following a single oral administration of aripiprazole (125 mg/kg), plasma and intrahepatic drug concentrations exceeded the in vitro EC_50_ but remained below the EC_90_ (Supplementary Figure 6D). Because the clinically used dose of aripiprazole in humans ranges from approximately 0.03 to 0.33 mg/kg/day, repeated administration at 10 mg/kg every 12 h was selected for pharmacokinetic simulation and in vivo efficacy studies ^18^. Pharmacokinetic simulation using the previously reported elimination half-life (4.1 h) ^19^ predicted that repeated administration (10 mg/kg every 12 h) would maintain plasma concentrations above the EC_50_ but below the EC_90_ (Supplementary Figure 6E). Stable HBV viremia was established in all transplanted mice with adequate human hepatocyte repopulation (Supplementary Figure S5A). Oral administration of aripiprazole (10 mg/kg every 12 h) was initiated at 32 days post-inoculation, resulting in a significant reduction in serum HBV DNA levels after two weeks compared with untreated controls (Supplementary Figure S5B and Figure 6F).

Immunohistochemical analysis revealed reduced intrahepatic expression of HBsAg and HBcAg in mice treated with vorapaxar or aripiprazole (Supplementary Figure S6). Neither compound affected body weight, serum cholinesterase levels, liver histology, or serum biochemical parameters (Supplementary Figures S6 and S7), indicating good tolerability at the tested doses. These findings demonstrate that vorapaxar and aripiprazole suppress HBV replication in vivo without detectable toxicity.

## Discussions

In the present study, we identified vorapaxar and aripiprazole as novel anti-HBV candidates by screening 1,470 FDA-approved compounds using an EnhI/Xp-driven reporter system. Both compounds suppressed HBV replication in multiple in vitro models, including HBV-replicating cells, HBV-infected HepG2-hNTCP cells, and primary human hepatocytes (Supplementary Figures S1 and S2). Mechanistically, aripiprazole suppressed CURS-BCP activity through ERK/JNK-dependent downregulation of HNF4α, whereas vorapaxar inhibited CURS-BCP independently of ERK/JNK signaling (Figure 1). Both compounds also inhibited EnhI/Xp activity and reduced STAT3 phosphorylation, although through distinct upstream pathways (Figures 2 and 3). Vorapaxar suppressed PAR-1-mediated SRC, EGFR, and STAT3 signaling, whereas aripiprazole inhibited SRC and STAT3 activation without affecting EGFR phosphorylation (Figures 3 and 4). In addition, PAR-1 activation enhanced HBV promoter activity and viral replication, whereas PAR-1 knockdown exerted the opposite effects (Figures 4 and 5), demonstrating that PAR-1–STAT3 signaling positively regulates HBV transcription. Finally, both vorapaxar and aripiprazole exhibited antiviral activity in human liver chimeric mice (Supplementary Figures S4 to S6 and Figures 6). Collectively, these findings identify PAR-1 signaling as a previously unrecognized regulator of HBV transcription while further supporting the established role of STAT3 in HBV replication. They also suggest that pharmacological inhibition of host signaling pathways controlling viral transcription represents a promising therapeutic strategy for chronic hepatitis B. Because these compounds target host pathways rather than viral proteins, their antiviral effects are expected to be maintained irrespective of viral sequence variation and may suppress HBV transcription from both cccDNA and integrated HBV DNA.

Deletion and reporter analyses identified D4 and D5 within enhancer I as the principal regions targeted by both vorapaxar and aripiprazole. These domains contain putative STAT3-binding sites, and both compounds consistently reduced STAT3 phosphorylation. Previous studies have demonstrated that STAT3 positively regulates HBV transcription and replication ^7^, that EGFR promotes HBV replication through STAT3 phosphorylation ^10^, and that phosphorylated STAT3 directly binds enhancer I on HBV cccDNA to enhance viral transcription through the HGF–MET–STAT3 signaling pathway ^20^. Furthermore, inhibition of JAK2/STAT3 signaling by cucurbitacin I suppresses HBV replication and destabilizes cccDNA ^21^. In addition, FOXA, whose putative binding site is located within D5 (Supplementary Figure S3), positively regulates enhancer I activity and cooperates with STAT3 ^7^. Together, these findings support the notion that vorapaxar and aripiprazole suppress enhancer I activity through inhibition of STAT3-dependent transcription, both directly through STAT3-responsive elements within D4 and indirectly through FOXA-associated transcriptional regulation. An important finding of this study is the identification of PAR-1 signaling as a previously unrecognized regulator of HBV transcription. PAR-1 knockdown reduced EnhI/Xp activity, CURS-BCP activity, and HBV RNA levels, whereas activation of PAR-1 with a tethered-ligand peptide produced the opposite effects. In addition, PAR-1 expression positively correlated with activation of the SRC–EGFR–STAT3 signaling axis. These findings support a model in which PAR-1 promotes HBV transcription through activation of signaling pathways converging on enhancer I/X promoter activity. To our knowledge, this is the first study demonstrating that PAR-1 signaling regulates HBV gene expression and replication. Although vorapaxar is a well-characterized PAR-1 antagonist, aripiprazole produced similar downstream effects despite having no known activity against PAR-1. Aripiprazole has been reported to directly inhibit SRC kinase activity ^22^, consistent with our observation that it reduced SRC and STAT3 phosphorylation without affecting EGFR phosphorylation (Figure 3). These findings suggest that vorapaxar and aripiprazole suppress a common STAT3-dependent transcriptional network through distinct upstream signaling pathways. Identification of the direct molecular target responsible for aripiprazole-mediated STAT3 inhibition will be an important subject for future investigation.

An important finding of this study is that both vorapaxar and aripiprazole suppressed not only EnhI/Xp activity but also CURS-BCP activity, although through distinct molecular mechanisms (Figure 1). Aripiprazole reduced HNF4α protein levels through an ERK/JNK-dependent pathway. Treatment with MG132 or ERK/JNK inhibitors restored both HNF4α protein expression and CURS-BCP activity, whereas neither treatment rescued EnhI/Xp activity in aripiprazole-treated cells (Figure 1). These findings indicate that HNF4α downregulation contributes to the inhibitory effect of aripiprazole on CURS-BCP activity but is dispensable for suppression of EnhI/Xp activity. Previous studies have shown that ERK activation suppresses HBV promoter activity through downregulation of HNF4α ^23^ and that ERK-mediated phosphorylation reduces the chromatin-binding activity of HNF4α ^24^. Consistent with these findings, restoration of CURS-BCP activity by ERK inhibition in the present study was accompanied by recovery of HNF4α protein levels. We previously demonstrated that TRIM21 promotes HNF4α degradation ^8^. Although a direct link between ERK/JNK signaling and TRIM21 induction has not been established, inflammatory signaling downstream of ERK/JNK may contribute to TRIM21 expression ^25^. Further studies will therefore be required to clarify the molecular mechanism underlying aripiprazole-induced HNF4α downregulation. In contrast, vorapaxar suppressed CURS-BCP activity without altering HNF4α protein levels or ERK/JNK signaling (Figure 1), suggesting the involvement of an alternative regulatory mechanism. Because CURS-BCP contains FOXA-binding sites ^26^ and PAR-1 signaling regulates STAT3 activation, STAT3 may cooperate with FOXA to regulate CURS-BCP activity. Interestingly, activation of PAR family members has been reported to increase HNF4-related DNA-binding activity in inflamed tissues ^27^, raising the possibility that PAR-1 signaling influences HNF4α function under physiological conditions. Although vorapaxar did not alter HNF4α protein abundance in hepatoma cell lines, modulation of HNF4α activity rather than its expression may contribute to its antiviral activity in vivo. Further studies are needed to determine whether PAR-1 signaling directly regulates HNF4α activity in hepatocytes and whether this contributes to HBV replication.

Aripiprazole has previously been reported to exhibit antiviral activity against several RNA viruses through mechanisms distinct from its antipsychotic effects. For example, it inhibits arenavirus infection by interfering with macropinocytosis-dependent viral entry ^28^ and suppresses SARS-CoV-2 pseudovirus entry through interaction with ACE2 ^29^. Although the molecular targets differ among viruses, these studies suggest that aripiprazole broadly modulates host pathways exploited during viral infection. In contrast to these previously reported mechanisms, the present study demonstrates that aripiprazole suppresses HBV replication primarily through inhibition of viral transcription. Specifically, aripiprazole reduced STAT3 phosphorylation and induced ERK/JNK-dependent downregulation of HNF4α, resulting in suppression of both EnhI/Xp and CURS-BCP activities. These findings extend the antiviral profile of aripiprazole by identifying a previously unrecognized host-targeting mechanism that suppresses HBV transcription and suggest that aripiprazole may have broad antiviral potential through modulation of distinct host signaling pathways.

Importantly, this study assessed the anti-HBV effects of vorapaxar and aripiprazole in vivo under dosing conditions intended to approximate their clinical use in humans. Both vorapaxar and aripiprazole exhibited antiviral activity in human liver chimeric mice at clinically relevant drug exposures without apparent severe toxicity (Supplementary Figures S8–6 and Figure 6). Because these compounds target host signaling pathways rather than viral proteins, their antiviral efficacy is expected to be maintained irrespective of viral sequence variation and may suppress HBV transcription from both cccDNA and integrated HBV DNA. These properties distinguish them from current nucleos(t)ide analogue therapies, which efficiently inhibit reverse transcription but have limited effects on viral transcription. Therefore, host-targeting transcriptional inhibitors such as vorapaxar and aripiprazole may complement existing antiviral therapies by targeting a distinct step in the HBV life cycle. Furthermore, optimization of these compounds or development of their derivatives may facilitate the clinical translation of transcription-targeting therapies for chronic hepatitis B.

## Acknowledgements

We thank C. Seeger and K. Watashi for providing plasmids and cell lines. We also thank M. Mori for her secretarial work and N. Yoneda, H. Kato, and T. Homma for their technical contribution to generating human liver chimeric mice. This work was supported by the Japan Science and Technology Agency (JST) Moonshot R&D under grant number JPMJMS2025, by the Japan Society for the Promotion of Science KAKENHI Grant Number JP25K02498, by Grants-in-Aid from the Japan Agency for Medical Research and Development (26fk0310526, 26fk0310533, 26fk0310545) and scholarship donations from Yakult Co., Ltd.

## Declaration of generative AI and AI-assisted technologies in the writing process

During the preparation of this work, the authors used ChatGPT (OpenAI) to improve the language, grammar, and readability of the manuscript. After using this tool, the authors carefully reviewed and edited the content as needed and take full responsibility for the content of the publication.

## Materials and Methods

### Ethics statement

All the experimental procedures and protocols used for this animal study were conducted in strict accordance with the Guide for the Care and Use of Laboratory Animals of the Central Institute for Experimental Medicine and Life Science (Permit Number: AIA260059).

### Cells, Viruses, and Compounds

The HepG2.2.15.7, HepG2-hNTCP-C4, and Hep38.7-Tet cell lines were kindly provided by K. Watashi, T. Wakita (National Institute of Infectious Diseases, Japan), and C. Seeger (Fox Chase Cancer Center) ^30^. HuH-7 cell line and derivatives were reported previously ^9^. The Huh7 GL4.18 CURS_BC_AeUS (CURS/BCP reporter) and Huh7 GL4.18 EnhI/X-Pro_AeUS (EnhI/Xp reporter) cell lines were established as described previously ^9,31^. HepG2 1.3×HBV-Luc and HepG2 CAG-HBV-Luc cells were described previously ^8^. HBV stocks for *in vitro* experiments were prepared from culture supernatants of Hep38.7-Tet cells as previously reported ^9,30,32^. HBV stock for *in vivo* experiments was prepared from HepG2.2.15 cells ^33^ and aliquoted into cryovial tubes and cryopreserved at −80 °C until use. Chimeric mice were intravenously inoculated with the same viral stock for all experiments.

An FDA-approved compound library consisting of 1,470 compounds was purchased from MedChemExpress (Monmouth Junction, NJ, USA). Compounds were supplied as DMSO stock solutions and diluted in culture medium immediately before use. TFLLR-NH2 was purchased from Selleck (CAS No.197794-83-5).

### Human Liver Chimeric Mice

Female F1 hybrid NOG-TKm30 mice, which are NOG mice expressing HSV-tk mutant clone 30 under the control of the mouse transthyretin gene enhancer/promoter, were used as recipients for primary human hepatocyte (PHH) transplantation^34^. Human liver chimeric mice were generated by transplanting 1 million cells/mouse of PHHs into the spleen of NOG-TKm30 mice with liver injury induced by oral administration of gancyclovir. The degree of human hepatocyte repopulation was assessed by measuring plasma cholinesterase (ChE) levels. Mice with stable and sufficient human hepatocyte engraftment were used for subsequent experiments.

Other materials and methods are described in Supplementary information.

## Supplementary information

### Supplementary Materials and Methods

#### Reporter Assays and Compound Screening

CURS/BCP and EnhI/Xp reporter cells were seeded in 48-well plates and treated with the indicated concentrations of vorapaxar or aripiprazole. Firefly luciferase (Fluc) activity and cell viability were determined as described previously ^1^. For primary screening, EnhI/Xp reporter cells were treated with each compound at 20 μM for 24 h. Compounds reducing Fluc activity to <35% of vehicle-treated controls while maintaining cell viability >80% were considered hit compounds. The resulting 22 hits were further evaluated using serial two-fold dilutions, and EC50 and CC50 values were calculated from dose–response curves to identify more effective compounds.

#### Plasmid Construction and Transient Transfection

Reporter plasmids containing the EnhI/Xp region lacking each of the D1–D7 domains (Supplementary Figure 1) were generated by inverse PCR using KOD FX Neo DNA polymerase (TOYOBO, Osaka, Japan) with the pGL4.18 EnhI/Xp reporter plasmid ^2^ as the template. The PCR products were digested with DpnI (New England Biolabs, Ipswich, MA, USA) at 37°C overnight to remove the template plasmid and subsequently circularized using the In-Fusion HD Cloning Kit (Takara Bio, Shiga, Japan). The resulting plasmids lacking the D1, D2, D3, D4, D5, D6, and D7 domains were designated pGL4.18EnhI-XproΔD1, pGL4.18EnhI-XproΔD2, pGL4.18EnhI-XproΔD3, pGL4.18EnhI-XproΔD4, pGL4.18EnhI-XproΔD5, pGL4.18EnhI-XproΔD6, and pGL4.18EnhI-XproΔD7, respectively. Primer sequences used for inverse PCR are listed in Supplementary Table S3

Six tandem copies of D3, D4, D5 or D6 were synthesized and cloned into pGL4.18[luc2P/Neo] (Promega, Madison, WI, USA) by GenScript (Piscataway, NJ, USA). The resulting plasmids including six tandem copies of D3, D4, D5, D6 and D7 were designated as pGL4.18EnhI-Xpro6XD3, pGL4.18EnhI-Xpro6XD4, pGL4.18EnhI-Xpro6XD5, and pGL4.18EnhI-Xpro6XD6, respectively. The synthetic DNA fragments consisted of six tandem repeats of the following sequences: D3, GGGCTTTGCTGCCCCATTTACACAATGTGGATATCCTGCC; D4, AGCTAAACAGGCTTTCACTTTCTCGCCAACTTACAAGGCCTTT; D5, GTACATGAACCTTTACCCCGTTGCTCGGCAACGGCCTG; and D6, TCTGTGCCAAGTGTTTGCTGACGCAACCCCCACT. PAR-1 cDNA was amplified using total RNA prepared from Huh-7 cells and introduced into pEF FLAG Gs pGKpuro ^3^. The resulting plasmid was used as PAR-1 expression plasmid.

For promoter assays, HEK293T/17 cells were co-transfected with either Fluc promoter reporter plasmids together with pRL-TK and pCAG-bsd-AP-HNF4α ^1^ or an empty plasmid. Activities of firefly luciferase (Fluc) and *Renilla* luciferase (Rluc) were measured using the Dual-Luciferase Reporter Assay System (Promega), and Fluc activity was normalized to Rluc activity.

#### Immunoblotting and RNA Interference

Cell lysis, immunoprecipitation, and immunoblotting were performed as previously described ^3^. Proteins were separated on 5–20% SDS–PAGE gels (ATTO, Tokyo, Japan) and transferred to PVDF membranes. Immunoreactive bands were visualized using SuperSignal West Femto and a Fusion FX7S.EDGE imaging system.

PAR-1-specific siRNA (ID 4285) and non-targeting control siRNA were purchased from Thermo Scientific and introduced using Lipofectamine RNAiMAX. Reverse transfection was used for reporter cells, whereas forward transfection was used for HBV-infected HepG2-hNTCP-C4 and PXB cells.

#### HBV Infection Assays

HepG2-hNTCP-C4 cells and primary human hepatocytes (PXB cells; PhoenixBio, Hiroshima, Japan) were infected with HBV as described previously ^2^. PXB cells were harvested at 12 days post-infection. HBV DNA, cccDNA, and RNA levels were quantified by real-time PCR or RT-qPCR as described below. Cell viability was determined in parallel.

#### Quantitative PCR and RT-PCR

Viral DNA was extracted using the SMITEST EX-R&D kit (Medical and Biological Laboratories). Total RNA extraction and first-strand cDNA synthesis were performed as previously described ^3^. HBV DNA, cccDNA, HBV RNA, and host mRNA levels were quantified by real-time PCR or RT-qPCR ^1–3^. Expression levels were normalized to S18 rRNA.

PAR-1 and GAPDH expression was additionally analyzed by semi-quantitative RT-PCR. PCR products were confirmed as single bands of the expected size by agarose gel electrophoresis. Amplification products were analyzed after 25–40 cycles to ensure linear amplification.

#### Chromatin Immunoprecipitation and Flow Cytometry

Cell-surface PAR-1 expression was analyzed by flow cytometry. Cells were detached with 1 mM EDTA, stained with FITC-conjugated anti-human PAR-1 antibody on ice, and analyzed using a FACS Celesta flow cytometer (BD Life Sciences).

#### Pharmacokinetic analysis

Pharmacokinetic studies were conducted in male ICR mice (4 weeks old). The test compound (vorapaxar or aripiprazole) dissolved in 0.5% carboxymethylcellulose was administered as a single dose (125 mg/kg) via oral gavage. Blood samples were collected at 5 min, 0.5, 1, 2, 3, 4, and 5 or 6 h via cardiac puncture into EDTA tubes. Plasma was separated by centrifugation and stored at −20°C until analysis. Perfused liver was homogenized by PBS, and stored at −20°C until analysis. Compound concentrations were quantified by a validated HPLC method following protein precipitation with acetonitrile containing an internal standard. A one-compartment model with first-order absorption for the compound was fitted to the mean plasma concentration-time profile in mice after single oral administration, and then the plasma concentration after multiple administration was predicted.

#### Antiviral Compounds and Treatment Regimen

Aripiprazole and vorapaxar were administered via oral gavage according to their pharmacological properties. Vehicle-treated mice served as negative controls. Treatment duration ranged from 2 to 4 weeks.

#### Serum Virological Analysis

Serum samples were collected at baseline and at regular intervals during treatment. HBV DNA was isolated following the Qiamp MinElute Virus Spin Kit (Qiagen, Hilden, Germany). HBV DNA was eluted in 60 µl, and 5 µl was used per well in the HBV DNA qPCR reaction. HBV DNA levels were quantified by real-time PCR as described previously ^4^.

#### Immunohistochemistry

At the end of the experimental period, tissue samples were collected, fixed in 10% neutral-buffered formalin, and embedded in paraffin. Formalin-fixed, paraffin-embedded (FFPE) tissue sections (4–5 μm thick) were deparaffinized in xylene and rehydrated through a graded ethanol series. Antigen retrieval was performed by heat-induced epitope retrieval in citrate buffer (pH 6.0) or EDTA buffer (pH 9.0), followed by cooling to room temperature. Endogenous peroxidase activity was quenched by incubation with 3% hydrogen peroxide, and nonspecific binding was blocked using an appropriate blocking solution. Tissue sections were then incubated with primary antibodies at optimized dilutions, including a rabbit polyclonal anti-HBsAg antibody (Bio-Rad Antibodies, Oxford, UK; catalog no. OBT0990), a mouse monoclonal anti-HBcAg antibody (Anogen, Mississauga, ON, Canada; catalog no. HBM-021-5), and a human-specific keratin K8/K18 monoclonal antibody (Progen Biotechnik GmbH, Heidelberg, Germany; catalog no. 10502), either overnight at 4°C or for 60 min at room temperature. After washing, sections were incubated with horseradish peroxidase (HRP)-conjugated secondary antibodies or polymer-based detection reagents according to the manufacturer’s instructions. Immunoreactive signals were visualized using 3,3′-diaminobenzidine (DAB) as the chromogen, and sections were counterstained with hematoxylin, dehydrated, and mounted. Negative control sections were processed in parallel by omitting the primary antibody.

#### Mouse blood biochemistry

Blood was collected from mice via cardiac puncture into serum tubes, centrifuged, and serum was stored at −80°C until analysis. Biochemical parameters such as aspartate aminotransferase (AST), alanine aminotransferase (ALT), creatinine (Cre), lactate dehydrogenase (LDH), total bilirubin (T-Bil), gamma-glutamyl transferase (GGT), total cholesterol (T-Cho), total protein (T-Pro), creatine phosphokinase (CPK), and albumin (Alb), were measured using an automated clinical chemistry analyzer (Dri-Chem NX700, Fujifilm, Tokyo) with manufacturer-approved reagents and calibrators.

#### Statistical Analysis

Statistical analyses were performed using GraphPad Prism 10 (GraphPad Software, San Diego, CA, USA). Data are presented as mean ± standard deviation (SD). After confirming normality (Shapiro–Wilk test) and homogeneity of variance (F-test), comparisons between each treatment group and the corresponding control group were performed using Student’s t-test. A p-value < 0.05 was considered statistically significant (*p < 0.05; **p < 0.01).

---

**Supplementary Figure S1.**
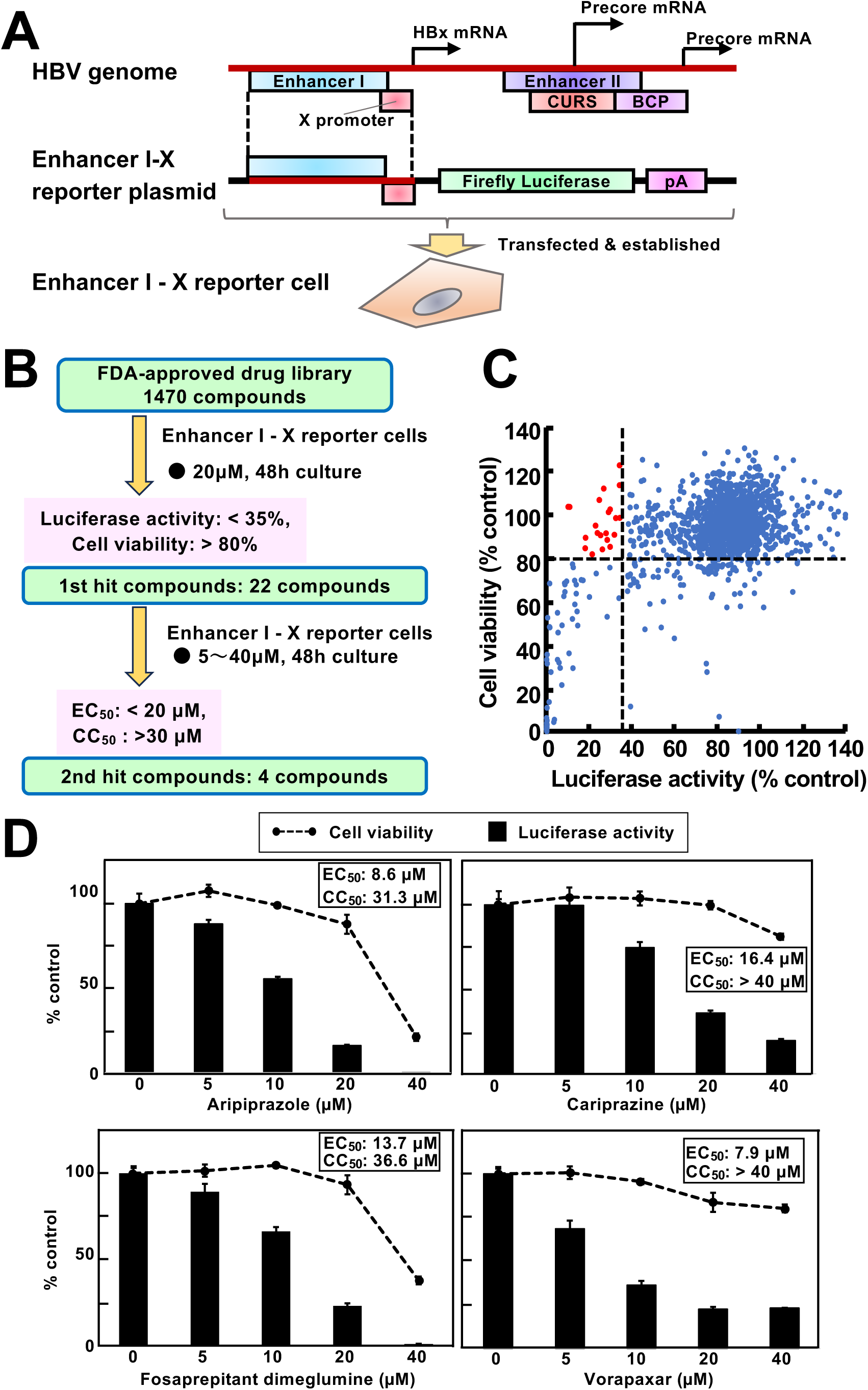
Identification of anti-HBV candidate compounds targeting EnhI/Xp. (A) Schematic representation of EnhI/Xp reporter cell line, including the reporter construct encoding luciferase under the control of EnhI/Xp. (B) Overview of the screening strategy using an FDA-approved drug library. Details of the screening procedure are described in the Materials and Methods. (C) Primary screening results showing the relationship between Enh I/Xp activity (luciferase activity) and cell viability. Each dot represents an individual compound. Red dots indicate hit compounds exhibiting less than 35% Fluc activity relative to vehicle-treated controls while maintaining greater than 80% cell viability. (D) Twenty-two compounds were identified as primary hits in the initial screening (C) and were subsequently evaluated for their EC₅₀ and CC₅₀ values using the Enh I/Xp reporter cells. Four compounds—aripiprazole, cariprazine, fosaprepitant dimeglumine, and vorapaxar—were selected as secondary hit compounds. Dose-dependent effects of these compounds on Enh I/Xp activity and cell viability are shown. Luciferase activity (bars) and cell viability (dashed lines) were measured in cells treated with each compound. The EC₅₀ and CC₅₀ values of aripiprazole, cariprazine, fosaprepitant dimeglumine, and vorapaxar are indicated in each graph.

**Supplementary Figure S2.**
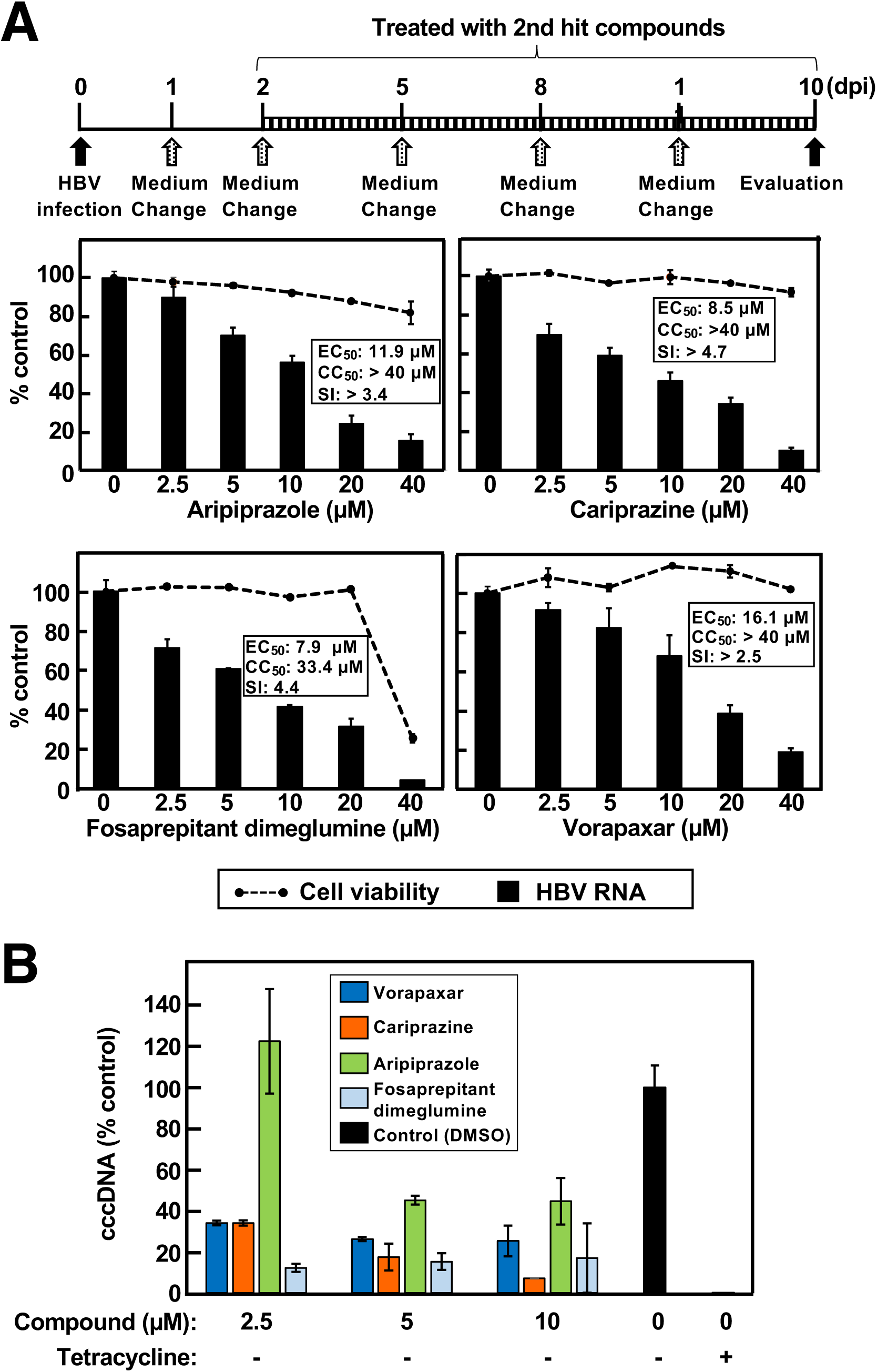
Inhibitory effects of hit compounds on HBV replication and cccDNA formation. (A) A schematic representation of the experimental timeline for HBV infection and treatment with the hit compounds (aripiprazole, cariprazine, fosaprepitant dimeglumine, and vorapaxar) is shown in the upper panel. PXB cells were infected with HBV at 5 genome equivalents (GEq)/cell and treated with the indicated concentrations of each compound. HBV DNA levels in the culture supernatants and cell viability were quantified as described in the Materials and Methods. (B) Hep38.7-Tet cells were cultured in the absence of tetracycline for 3 days and then further incubated for an additional 3 days without tetracycline in the presence of each compound. Cells were harvested 3 days after compound treatment and lysed. The amount of cccDNA was quantified as described in the Materials and Methods. Data are presented as the mean ± standard deviation (SD).

**Supplementary Figure S3.**
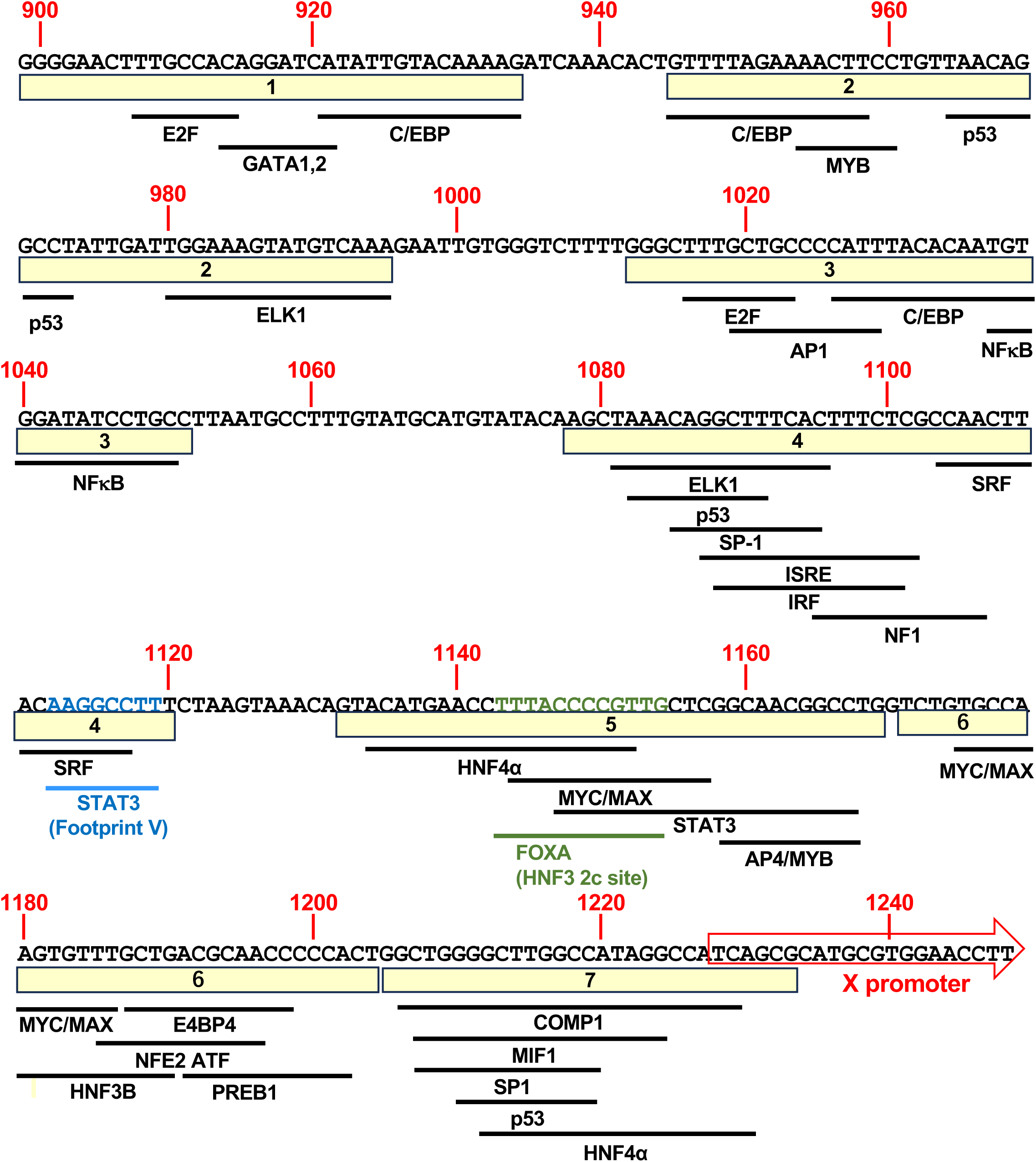
In silico prediction of transcription factor binding sites within the EnhI/Xp. The nucleotide sequence of EnhI/Xp was analyzed using the TFBIND program to predict potential transcription factor binding sites. Predicted binding sites are indicated below the DNA sequence. Based on the distribution of these sites, the EnhI/Xp region was subdivided into seven domains (D1–D7). These domains were subsequently used to construct the deletion mutants and six-tandem-repeat reporter plasmids analyzed in Figure 4.

**Supplementary Figure S4.**
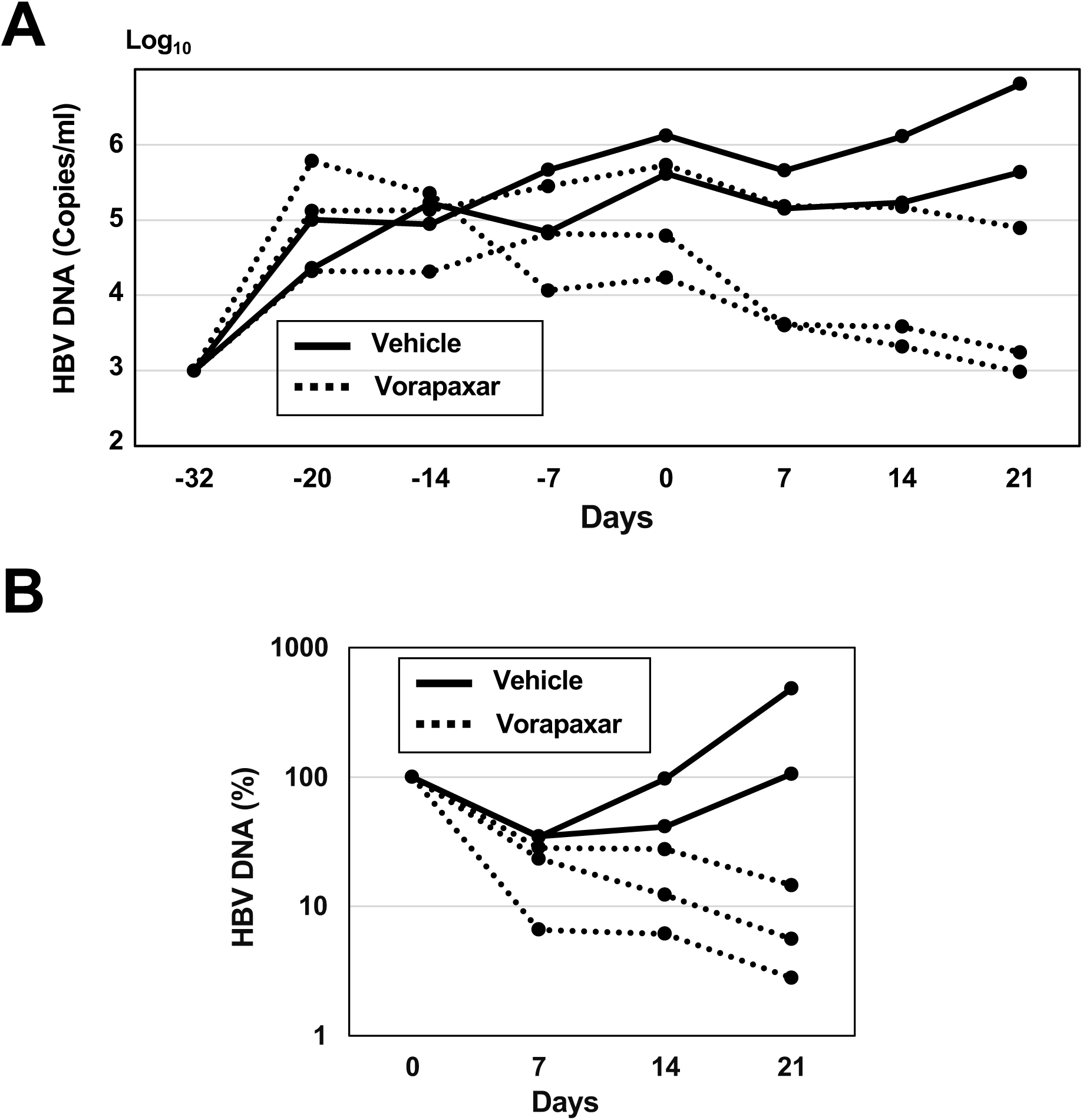
Antiviral effect of vorapaxar in HBV-infected NOG-TKm30 human liver chimeric mice. (A) Changes in serum HBV DNA levels in mice treated with vehicle or vorapaxar, corresponding to the experiment shown in Figure 6A–C. Mice with serum cholinesterase levels greater than 50 U/L, indicative of efficient and stable human hepatocyte engraftment, were inoculated with HBV (1 × 10^9 copies/mouse) via the tail vein on day −32. HBV-infected NOG-TKm30 human liver chimeric mice with stable viremia were orally administered vehicle or vorapaxar at a dose of 0.5 mg/kg twice daily for 21 consecutive days. (B) Individual time courses of serum HBV DNA levels, expressed as a percentage of baseline (Day 0 = 100%). Each line represents one mouse. Data are shown as mean ± SEM. Statistical significance was determined using appropriate parametric or non-parametric tests. A p value < 0.05 was considered statistically significant.

**Supplementary Figure S5.**
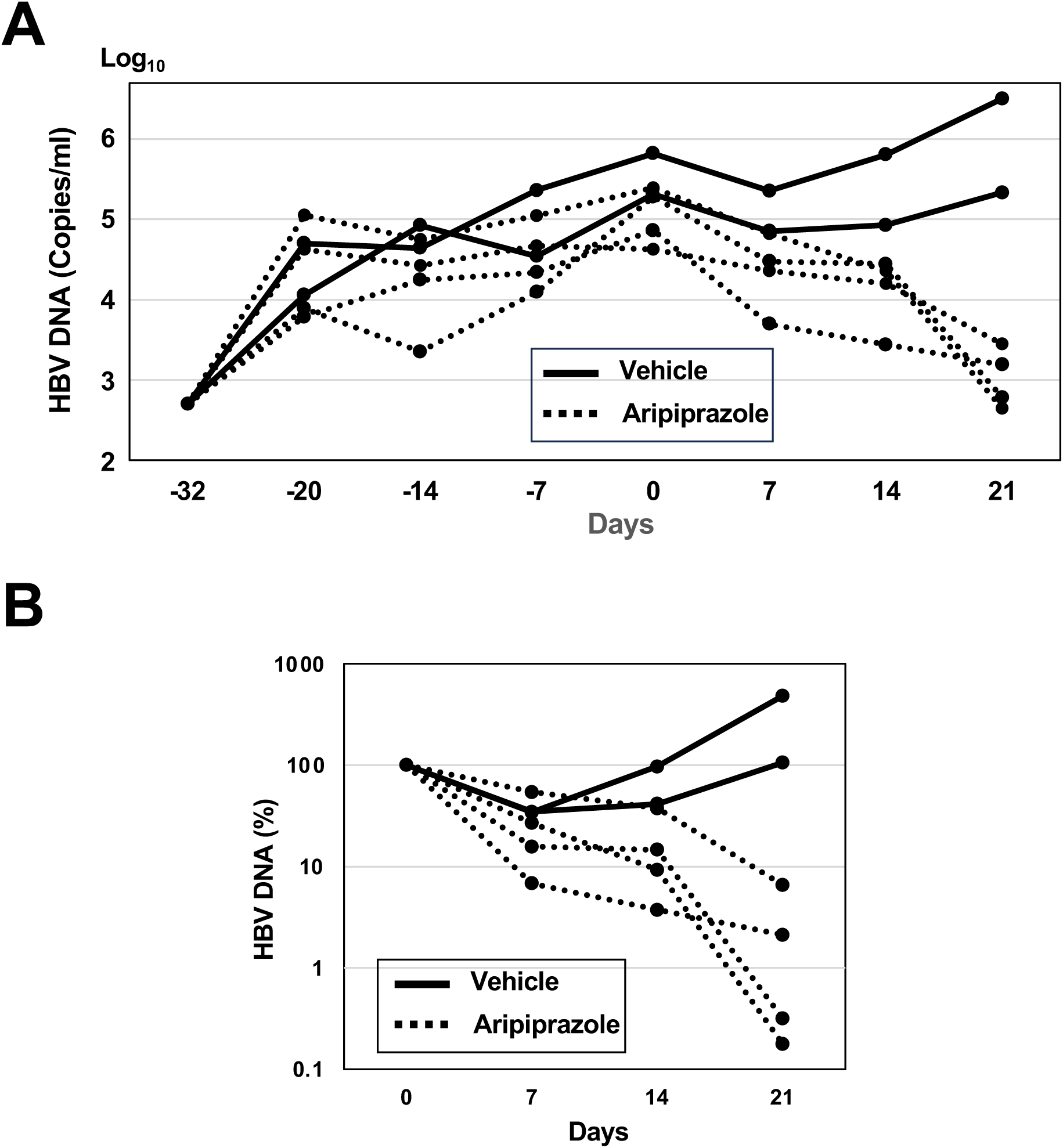
Antiviral effect of aripiprazole in HBV-infected NOG-TKm30 human liver chimeric mice. (A) Changes in serum HBV DNA levels in mice treated with vehicle or aripiprazole, corresponding to the experiment shown in Figure 6D–F. Mice with serum cholinesterase levels greater than 50 U/L, indicative of efficient and stable human hepatocyte engraftment, were inoculated with HBV (1 × 10^9 copies/mouse) via the tail vein on day −32. HBV-infected NOG-TKm30 human liver chimeric mice with stable viremia were orally administered vehicle or aripiprazole at a dose of 0.5 mg/kg twice daily for 21 consecutive days. (B) Individual time courses of serum HBV DNA levels, expressed as a percentage of baseline (Day 0 = 100%). Each line represents one mouse. Data are shown as mean ± SEM. Statistical significance was determined using appropriate parametric or non-parametric tests. A p value < 0.05 was considered statistically significant.

**Supplementary Figure S6.**
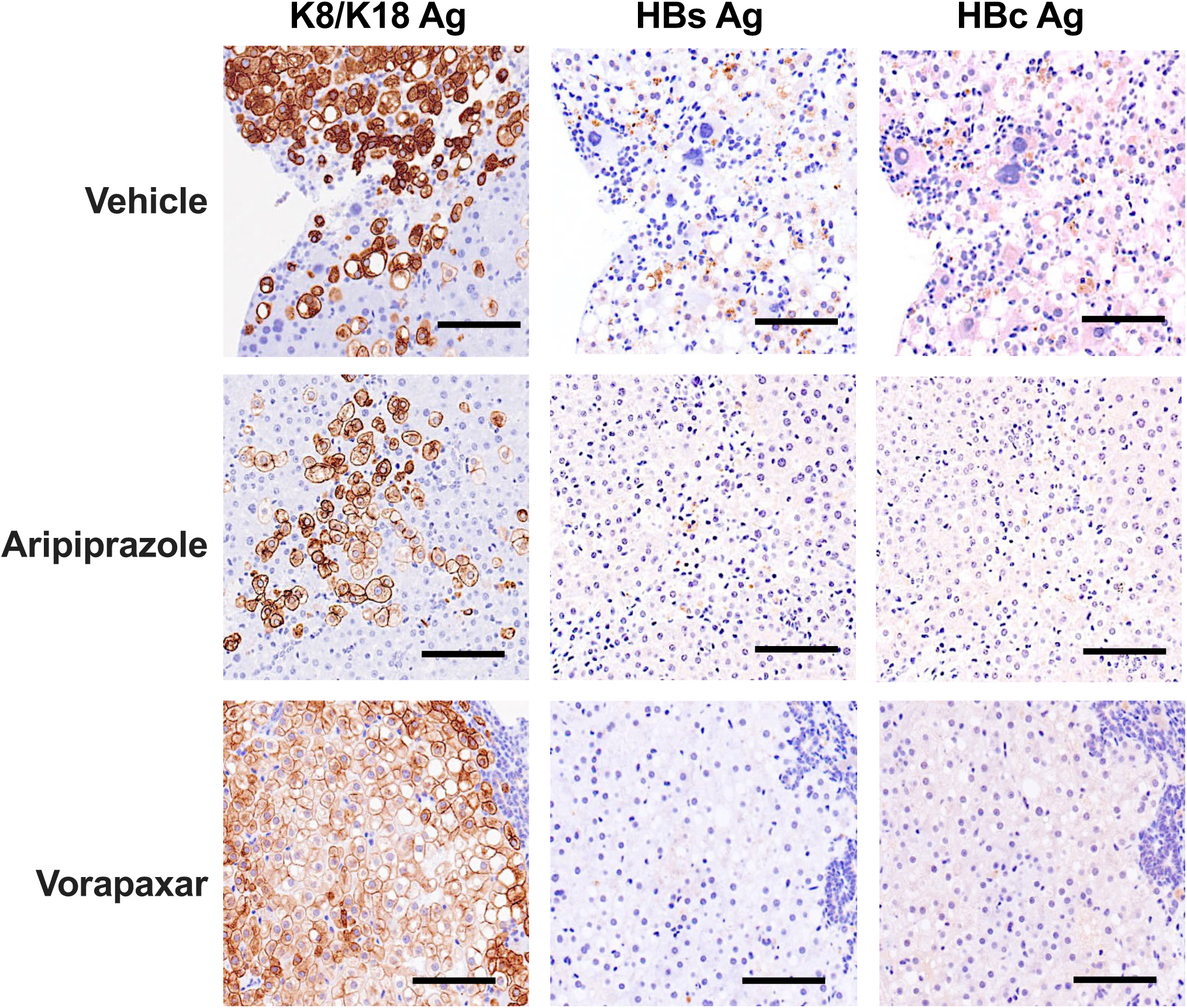
Immunohistochemical staining of HBsAg and HBcAg in liver tissues from sacrificed mice. Liver tissues were collected at the end of the experiment and subjected to immunohistochemical (IHC) analysis. Intrahepatic HBsAg and HBcAg were detected using a rabbit polyclonal anti-HBsAg antibody and a mouse monoclonal anti-HBcAg antibody, respectively. Signals were visualized with horseradish peroxidase (HRP)– conjugated secondary antibodies and 3,3′-diaminobenzidine (DAB) substrate (brown). Nuclei were counterstained with hematoxylin (blue). Human hepatocytes were identified by immunostaining with a human-specific keratin K8/K18 monoclonal antibody. Representative images from one mouse per group are shown. Scale bar, 100 μm.

**Supplementary Figure S7.**
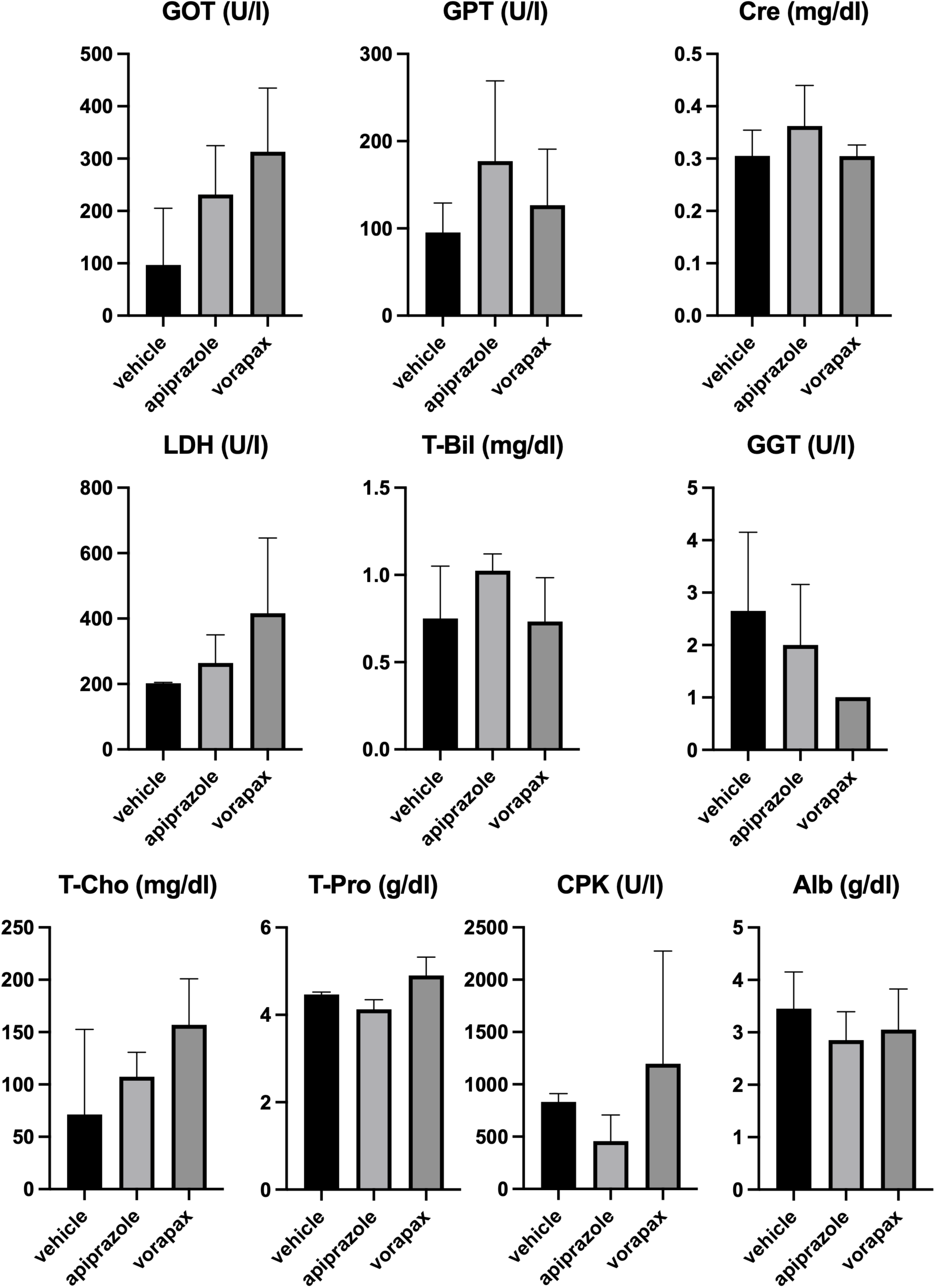
Safety profile and tolerability of the replication inhibitor. Evaluation of safety and tolerability of the replication inhibitors in NOG-TKm30 human liver chimeric mice. Biochemical parameters such as AST, ALT, Cre, LDH, T-Bil, GGT, T-Cho, T-Pro, CPK, or Alb were measured using an automated clinical chemistry analyzer. Data are presented as the mean ± SD. Homogeneity of variances was evaluated using the Brown–Forsythe test. One-way ANOVA followed by Dunnett’s multiple comparisons test was used when equal variances were confirmed. When the Brown–Forsythe test indicated unequal variances (P < 0.05), Brown–Forsythe and Welch ANOVA tests were used instead. Statistical analyses were performed using GraphPad Prism (version 10.6.0; GraphPad Software, San Diego, CA, USA). A two-sided P value < 0.05 was considered statistically significant.

**Supplementary Table S1.**
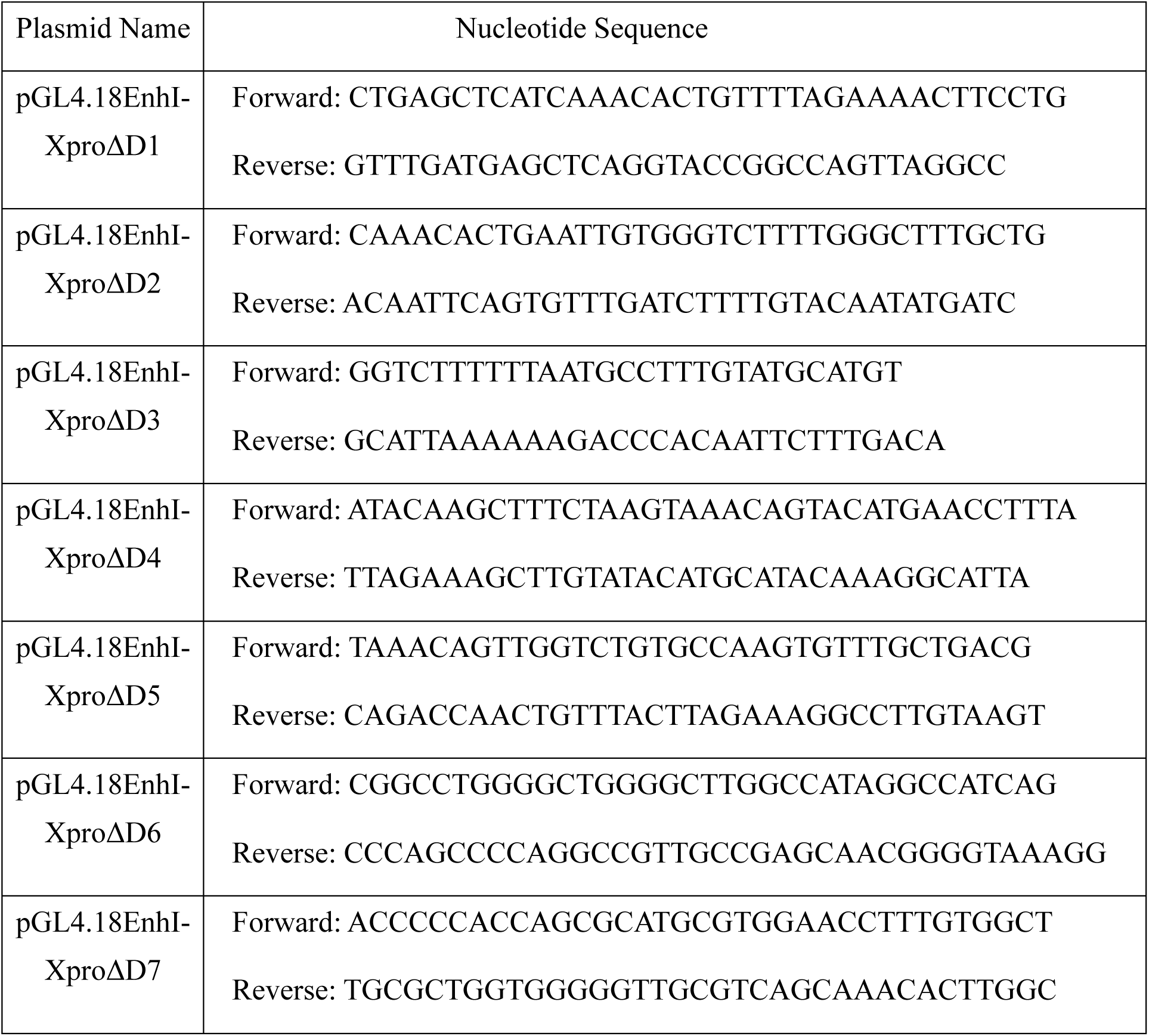
Primer pairs used for inverse PCR to construct reporter plasmids lacking D1-D7 domains.

## References

[1] Jeng WJ, Papatheodoridis GV, Lok ASF. Hepatitis B. Lancet 2023;401:1039–1052.

[2] Moolla N, Kew M, Arbuthnot P. Regulatory elements of hepatitis B virus transcription. J Viral Hepat 2002;9:323–331.

[3] Villanueva RA, Loyola A. The Intrinsically Disordered Region of HBx and Virus-Host Interactions: Uncovering New Therapeutic Approaches for HBV and Cancer. Int J Mol Sci 2025;26.

[4] Yu X, Mertz JE. Distinct modes of regulation of transcription of hepatitis B virus by the nuclear receptors HNF4alpha and COUP-TF1. J Virol 2003;77:2489–2499.

[5] Johnson JL, Raney AK, McLachlan A. Characterization of a functional hepatocyte nuclear factor 3 binding site in the hepatitis B virus nucleocapsid promoter. Virology 1995;208:147–158.

[6] Song Y, Zeng Y, Zheng H, et al. SOX6 is a novel host factor that promotes hepatitis B virus replication by enhancing the transcriptional activity of enhancer I. Antiviral Res 2026;247:106359.

[7] Waris G, Siddiqui A. Interaction between STAT-3 and HNF-3 leads to the activation of liver-specific hepatitis B virus enhancer 1 function. J Virol 2002;76:2721–2729.

[8] Yamashita A, Kasai H, Maekawa S, et al. Berberine promotes K(48)-linked polyubiquitination of HNF4alpha, leading to the inhibition of HBV replication. Antiviral Res 2024;232:106027.

[9] Yamashita A, Tamaki M, Kasai H, et al. Inhibitory effects of metachromin A on hepatitis B virus production via impairment of the viral promoter activity. Antiviral Res 2017;145:136–145.

[10] Gan CJ, Li WF, Li CN, et al. EGF receptor inhibitors comprehensively suppress hepatitis B virus by downregulation of STAT3 phosphorylation. Biochem Biophys Rep 2020;22:100763.

[11] Hosel M, Quasdorff M, Ringelhan M, et al. Hepatitis B Virus Activates Signal Transducer and Activator of Transcription 3 Supporting Hepatocyte Survival and Virus Replication. Cell Mol Gastroenterol Hepatol 2017;4:339–363.

[12] Bonaca MP, Morrow DA. SCH 530348: a novel oral thrombin receptor antagonist. Future Cardiol 2009;5:435–442.

[13] Morrison JT, Govsyeyev N, Hess CN, et al. Vorapaxar for Prevention of Major Adverse Cardiovascular and Limb Events in Peripheral Artery Disease. J Cardiovasc Pharmacol Ther 2022;27:10742484211056115.

[14] de Bartolomeis A, Tomasetti C, Iasevoli F. Update on the Mechanism of Action of Aripiprazole: Translational Insights into Antipsychotic Strategies Beyond Dopamine Receptor Antagonism. CNS Drugs 2015;29:773–799.

[15] Mori Y, Takeuchi H, Tsutsumi Y. Current perspectives on the epidemiology and burden of tardive dyskinesia: a focused review of the clinical situation in Japan. Ther Adv Psychopharmacol 2022;12:20451253221139608.

[16] Chackalamannil S, Wang Y, Greenlee WJ, et al. Discovery of a novel, orally active himbacine-based thrombin receptor antagonist (SCH 530348) with potent antiplatelet activity. J Med Chem 2008;51:3061–3064.

[17] Wang A. Review of vorapaxar for the prevention of atherothrombotic events. Expert Opin Pharmacother 2015;16:2509–2522.

[18] Luan S, Wan H, Zhang L, et al. Efficacy, acceptability, and safety of adjunctive aripiprazole in treatment-resistant depression: a meta-analysis of randomized controlled trials. Neuropsychiatr Dis Treat 2018;14:467–477.

[19] Nagasaka Y, Sano T, Oda K, et al. Impact of genetic deficiencies of P-glycoprotein and breast cancer resistance protein on pharmacokinetics of aripiprazole and dehydroaripiprazole. Xenobiotica 2014;44:926–932.

[20] Funato K, Miyake N, Sekiba K, et al. Cabozantinib inhibits HBV-RNA transcription by decreasing STAT3 binding to the enhancer region of cccDNA. Hepatol Commun 2023;7.

[21] Zhao L, Yuan H, Wang Y, et al. p-STAT3-elevated E3 ubiquitin ligase DTX4 confers the stability of HBV cccDNA by ubiquitinating APOBEC3B in liver. Theranostics 2024;14:6036–6052.

[22] Kim MS, Yoo BC, Yang WS, et al. Src is the primary target of aripiprazole, an atypical antipsychotic drug, in its anti-tumor action. Oncotarget 2018;9:5979–5992.

[23] Dai XQ, Cai WT, Wu X, et al. Protocatechuic acid inhibits hepatitis B virus replication by activating ERK1/2 pathway and down-regulating HNF4alpha and HNF1alpha in vitro. Life Sci 2017;180:68–74.

[24] Veto B, Bojcsuk D, Bacquet C, et al. The transcriptional activity of hepatocyte nuclear factor 4 alpha is inhibited via phosphorylation by ERK1/2. PLoS One 2017;12:e0172020.

[25] Nikolaou KC, Godbersen S, Manoharan M, et al. Inflammation-induced TRIM21 represses hepatic steatosis by promoting the ubiquitination of lipogenic regulators. JCI Insight 2023;8.

[26] Gerner P, Lausch E, Friedt M, et al. Hepatitis B virus core promoter mutations in children with multiple anti-HBe/HBeAg reactivations result in enhanced promoter activity. J Med Virol 1999;59:415–423.

[27] Saban R, Simpson C, Davis CA, et al. Transcription factor network downstream of protease activated receptors (PARs) modulating mouse bladder inflammation. BMC Immunol 2007;8:17.

[28] Herring S, Oda JM, Wagoner J, et al. Inhibition of Arenaviruses by Combinations of Orally Available Approved Drugs. Antimicrob Agents Chemother 2021;65.

[29] Lu J, Hou Y, Ge S, et al. Screened antipsychotic drugs inhibit SARS-CoV-2 binding with ACE2 in vitro. Life Sci 2021;266:118889.

[30] Ogura N, Watashi K, Noguchi T, et al. Formation of covalently closed circular DNA in Hep38.7-Tet cells, a tetracycline inducible hepatitis B virus expression cell line. Biochem Biophys Res Commun 2014;452:315–321.

[31] Yamashita A, Fujimoto Y, Tamaki M, et al. Identification of Antiviral Agents Targeting Hepatitis B Virus Promoter from Extracts of Indonesian Marine Organisms by a Novel Cell-Based Screening Assay. Mar Drugs 2015;13:6759–6773.

[32] Ladner SK, Otto MJ, Barker CS, et al. Inducible expression of human hepatitis B virus (HBV) in stably transfected hepatoblastoma cells: a novel system for screening potential inhibitors of HBV replication. Antimicrob Agents Chemother 1997;41:1715–1720.

[33] Sells MA, Chen ML, Acs G. Production of hepatitis B virus particles in Hep G2 cells transfected with cloned hepatitis B virus DNA. Proc Natl Acad Sci U S A 1987;84:1005–1009.

[34] Uehara S, Higuchi Y, Yoneda N, et al. An improved TK-NOG mouse as a novel platform for humanized liver that overcomes limitations in both male and female animals. Drug Metab Pharmacokinet 2022;42:100410.

## References

[1] Yamashita A, Kasai H, Maekawa S, et al. Berberine promotes K(48)-linked polyubiquitination of HNF4alpha, leading to the inhibition of HBV replication. Antiviral Res 2024;232:106027.

[2] Yamashita A, Tamaki M, Kasai H, et al. Inhibitory effects of metachromin A on hepatitis B virus production via impairment of the viral promoter activity. Antiviral Res 2017;145:136–145.

[3] Kasai H, Yamashita A, Akaike Y, et al. HCV infection activates the proteasome via PA28gamma acetylation and heptamerization to facilitate the degradation of RNF2, a catalytic component of polycomb repressive complex 1. mBio 2024;15:e0169124.

[4] Fukano K, Oshima M, Tsukuda S, et al. NTCP Oligomerization Occurs Downstream of the NTCP-EGFR Interaction during Hepatitis B Virus Internalization. J Virol 2021;95:e0093821.

